# A nonlinear code for event probability in the human brain

**DOI:** 10.1101/2024.02.28.582455

**Authors:** Cedric Foucault, Tiffany Bounmy, Sébastien Demortain, Bertrand Thiririon, Evelyn Eger, Florent Meyniel

## Abstract

Assessing probabilities and predicting future events are fundamental for perception and adaptive behavior, yet the neural representations of probability remain elusive. While previous studies have shown that neural activity in several brain regions correlates with probability-related factors such as surprise and uncertainty, similar correlations have not been found for probability. Here, using 7 Tesla functional magnetic resonance imaging, we uncover a representation of the probability of the next event in a sequence within the human dorsolateral prefrontal and intraparietal cortices. Crucially, univariate and multivariate analyses revealed that this representation employs a highly nonlinear code. Tuning curves for probability exhibit selectivity to various probability ranges, while the code for confidence accompanying these estimates is predominantly linear. The diversity of tuning curves we found recommends that future studies move from assuming linear correlates or simple canonical forms of tuning curves to considering richer representations whose benefits remain to be discovered.

## Introduction

Our world is conveniently described in terms of probabilities. Examples include perception (e.g. “I’m sure I saw someone I know in the crowd” (Kersten et al., 2004)), social interaction (“My boss is probably lying about my pay rise” (Diaconescu et al., 2014)), and prediction (“It is unlikely to rain tomorrow” (Yang & Shadlen, 2007)). These probabilities reflect the uncertainty of our beliefs about the world, due to the ambiguity of our inputs or limitations in our information processing capabilities (Walker et al., 2023). Human behavior is adapted to these probabilities, as shown by numerous examples both in laboratory experiments and in real life. To name a few, perception is tuned to the probability of occurrence of a given object or feature in the visual, auditory, and somatosensory spaces (Summerfield & de Lange, 2014), and choices are guided by the probability of reward (Rangel et al., 2008). These behavioral results imply that probabilities must be encoded in the brain.

However, the neural code for probabilities remains largely unknown. Several studies have reported that neural activity increases (or decreases) monotonically with quantities that are related to probabilities, but not with probabilities themselves. Examples of related quantities include the coding of prediction error (Schultz et al., 1997), surprise (O’Reilly et al., 2013), and predictability (Bach et al., 2011; Fiorillo, 2003; Monosov & Hikosaka, 2013). Some studies explicitly reported a failure to identify correlations between probabilities and neural activity (Lebreton et al., 2015; Marshall et al., 2022).

Here, we tested the possibility that neural activity does not scale with probability levels, but instead that the neural code for probability is nonlinear. The codes we studied belong to the broader class of codes that are characterized by tuning curves. A tuning curve relates the value of a quantitative feature (e.g. the orientation of a grating (Hubel & Wiesel, 1959), the number of items in a set (Eger, 2016; Moser et al., 2008; Nieder, 2016)) to the neural activity that this value elicits on average across trials. A special case of tuning curve is the linear code, where neural activity is simply proportional to the encoded quantity. The linear code has been extensively tested and it accounts for the encoding of various quantities such as prediction error, surprise, predictability as mentioned above, and other quantities not related to probabilities, e.g. value (Padoa-Schioppa & Assad, 2008), complexity (Wang et al., 2019), salience (Kutlu et al., 2021). There are also numerous examples of quantities that are encoded with a nonlinear code, such as orientation (Hubel & Wiesel, 1959), numerosity (Nieder, 2016), proportion (Jacob & Nieder, 2009). For these quantities, tuning curves are not linear, but highly non-monotonic. They are often bell-shaped, but more complex tuning curves have also been observed (Diehl et al., 2017; Hardcastle et al., 2017).

To adjudicate between linear and nonlinear codes for probability, it is necessary to characterize the brain’s tuning curves for probability. We measured neural activity with ultra-high field fMRI (7T) because it provides a high signal-to-noise ratio to capture the organization of the neural code at a millimeter-scale with a whole-brain coverage (which is very useful given the lack of anatomical priors on the brain regions involved in the representation of probabilities). We characterized tuning curves for probability in each fMRI voxel using a method that combines the generality of function approximation with basis functions (Bishop, 2007) and linearizing encoding models (Huth et al., 2016; Naselaris et al., 2011). This method does not make assumptions about tuning shape, in contrast to related encoding methods (see Discussion, (Dumoulin & Wandell, 2008; Friston et al., 2007)). It is also more informative about tuning curves than decoding methods. For example, linear classifiers (e.g. regularized linear regression (Findling et al., 2023), support vector machine (Eger et al., 2009)) are often used to decode a quantity from brain signals, but these methods can achieve high performance regardless of whether the underlying tuning curves are linear, bell-shaped, or even more complex (Kriegeskorte & Diedrichsen, 2019). We analyzed the tuning curves for probability both at the single-voxel level and at the level of populations of voxels to strengthen our conclusions. At the single-voxel level, following a univariate approach, we quantified the form and degree of nonlinearity of tuning curves. At the population level, following a multivariate approach, we quantified whether patterns of voxel responses (more specifically their similarity across probabilities) conformed to a linear or a highly nonlinear code.

In terms of experimental design, studying the neural representation of probability requires minimizing potential confounds between probability, and other constructs and task features (Walker et al., 2023). For example, in several previous studies the concept of probabilities has been confounded with evidence for a decision (Yang & Shadlen, 2007), or with the reward that the subject expects to receive (Ferrari-Toniolo & Schultz, 2023; Kepecs et al., 2008). Probabilities are also correlated with some motor or sensorimotor transformations in many tasks (Yang & Shadlen, 2007), which is useful for studying the behavioral relevance of probabilities, but is a nuisance when studying the neural code of probabilities themselves. To alleviate the problem of confounding factors and variables, we used a probability learning paradigm. Participants estimated the (changing) generative probability of occurrence of neutral items presented sequentially. To limit confounds related to motor actions and decision making, this estimation was covert on most trials (on which the analysis focused), and behavioral reports of probabilities were kept to a minimum. The design had no overt rewards to avoid confounds related to valuation processes. Finally, probability is a latent parameter of this task (i.e., it is not observable in stimulus space), which minimizes confounds related to sensory processes.

To anticipate our result, we provide evidence for the nonlinearity of the neural code of probability, and we strengthen this conclusion by comparing it to another quantity whose code is linear. There is evidence from previous studies that the confidence that accompanies a probability estimate in a learning task correlates (linearly) with neural activity, particularly in regions of the fronto-parietal network (Bounmy et al., 2023; McGuire et al., 2014; Meyniel & Dehaene, 2017; Tomov et al., 2020). The comparison between probability and confidence is not confounded by between-subject differences, or differences in perceptual space since both are derived from the very same sequence of stimuli.

## Results

### Task probing the representation of probabilities

We subjected twenty-six human subjects to a probability learning task and simultaneously measured their brain activity using ultra-high field (7T) fMRI, to examine their neural representation of probabilities, and of the confidence associated with their probability estimates. After one training session out of the scanner, subjects performed four sessions of the task in the MRI scanner. In each session, subjects observe a sequence of 420 stimuli appearing one by one, each with a binary value (A or B) sampled from a hidden generative probability p(A) (Fig. 1). This hidden probability undergoes abrupt change points at random unpredictable times which were not signaled to the subjects. The subject’s goal is to estimate the current value of the hidden probability throughout the sequence. To perform the task correctly, subjects have to frequently update their estimate as new observations are made. This experimental protocol allows us to efficiently probe the neural representation of probabilities and confidence as it induces frequent variations in these quantities that can be compared with variations in measured neural activity.

**Figure 1.**
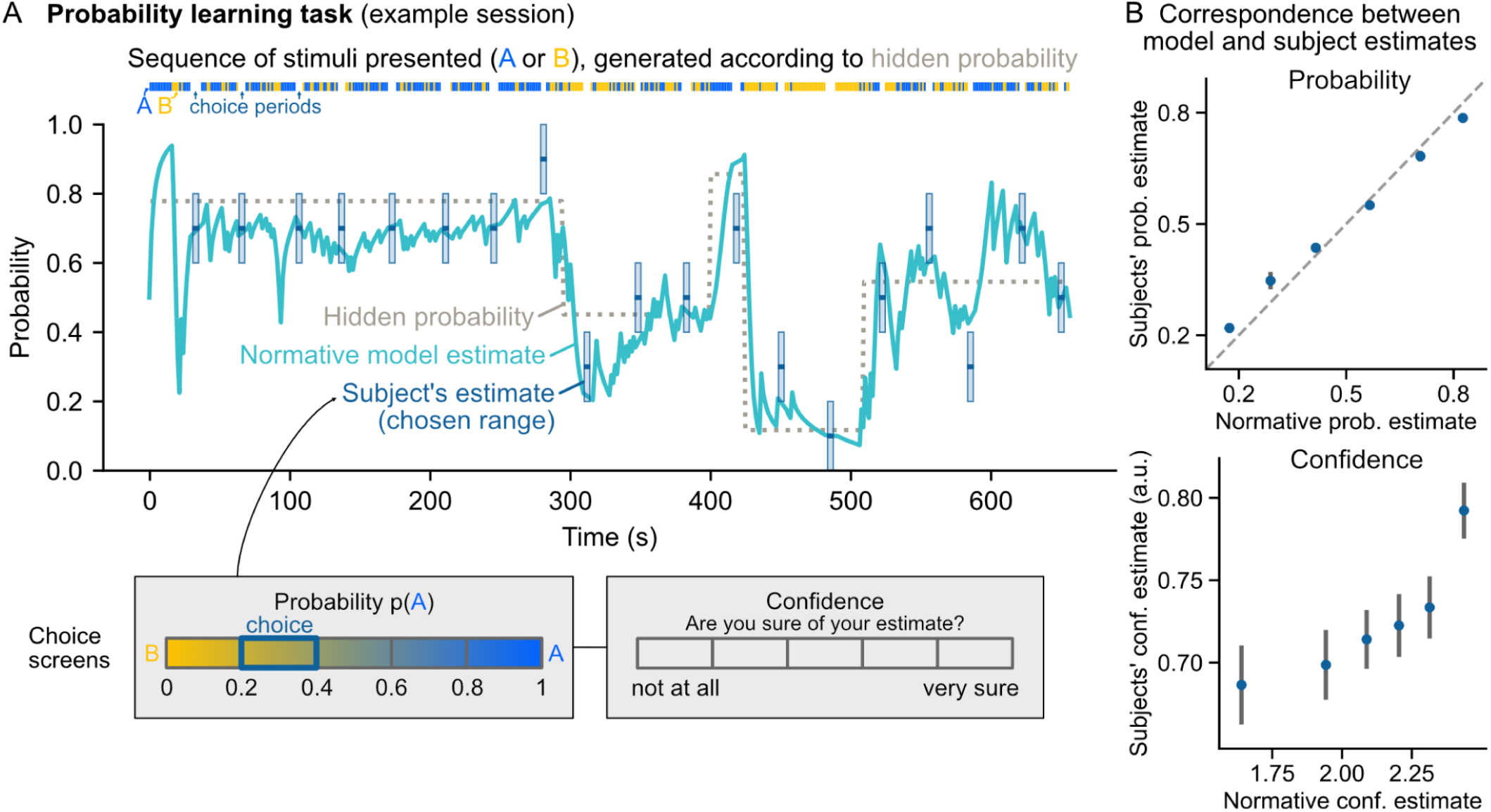
Task and variables of interest. (A) Task session. Visual stimuli of two types, A or B, generated according to a hidden probability p(A), are presented successively to the subject. During the sequence, the hidden probability changes at random and unpredictable times. Throughout the session, the subject must estimate the current value of the hidden probability. Occasionally between two stimuli, a choice period occurs during which the subject chooses a range for their current probability estimate and a confidence level associated with their estimate. The normative model of the task is used to obtain the trial-by-trial estimates that the subject should neurally represent. The values of these different variables are displayed above for an example session. (B) Behavioral results: The estimates chosen by the subjects and the normative estimates are very close for probability (top), and quite correlated for confidence (bottom). Subjects’ estimates were binned into six equal quantiles of normative estimate, averaged at the subject level and then at the group level. Subjects’ confidence was recorded from 0 to 1; normative confidence is in log precision units (hence the different scales). Dots and error bars show group-level mean±s.e.m. Dashed diagonal is the identity line.

To minimize disruption to the fMRI signals, we asked subjects to perform their estimation covertly and only occasionally requested behavioral reports. The reports occurred during dedicated periods (on average every 22 stimuli), during which the subject had to choose a range for their current probability estimate and associate a level of confidence with that estimate. We found a strong correspondence between the estimates reported by the subjects and those of the normative model (see Methods) (Fig. 1 B; Pearson *r*=0.79 mean ±0.12 s.e.m, *t_25_*=31.8, p<10^-21^ for probability and *r*=0.16±0.13, *t_25_*=5.9, p<10^-5^ for confidence). In another recent study, we found that this strong correspondence with the normative model was consistently observed at the trial-by-trial level (Foucault & Meyniel, 2023). In what follows, we thus used the estimates of the normative model as a proxy for those of the subjects in order to relate trial-by-trial estimates with ongoing neural activity.

### Encoding models and evaluation procedure

To relate probability and confidence estimates to neural activity, we constructed models that predict neural activity as a function of the estimate of interest (probability or confidence), which we refer to as encoding models (Fig. 2A). We defined two classes of models: the class of linear encoding models, which is the one classically used in fMRI studies and which assumes a linear relationship between the variable of interest and neural activity, and the class of versatile encoding models, which does not make such an assumption. Instead, the versatile class captures all kinds of relationships, which include but are not restricted to linear relationships, by leveraging approximation theory. The versatile class can approximate arbitrary functions as a weighted sum of basis functions (see Methods).

**Figure 2.**
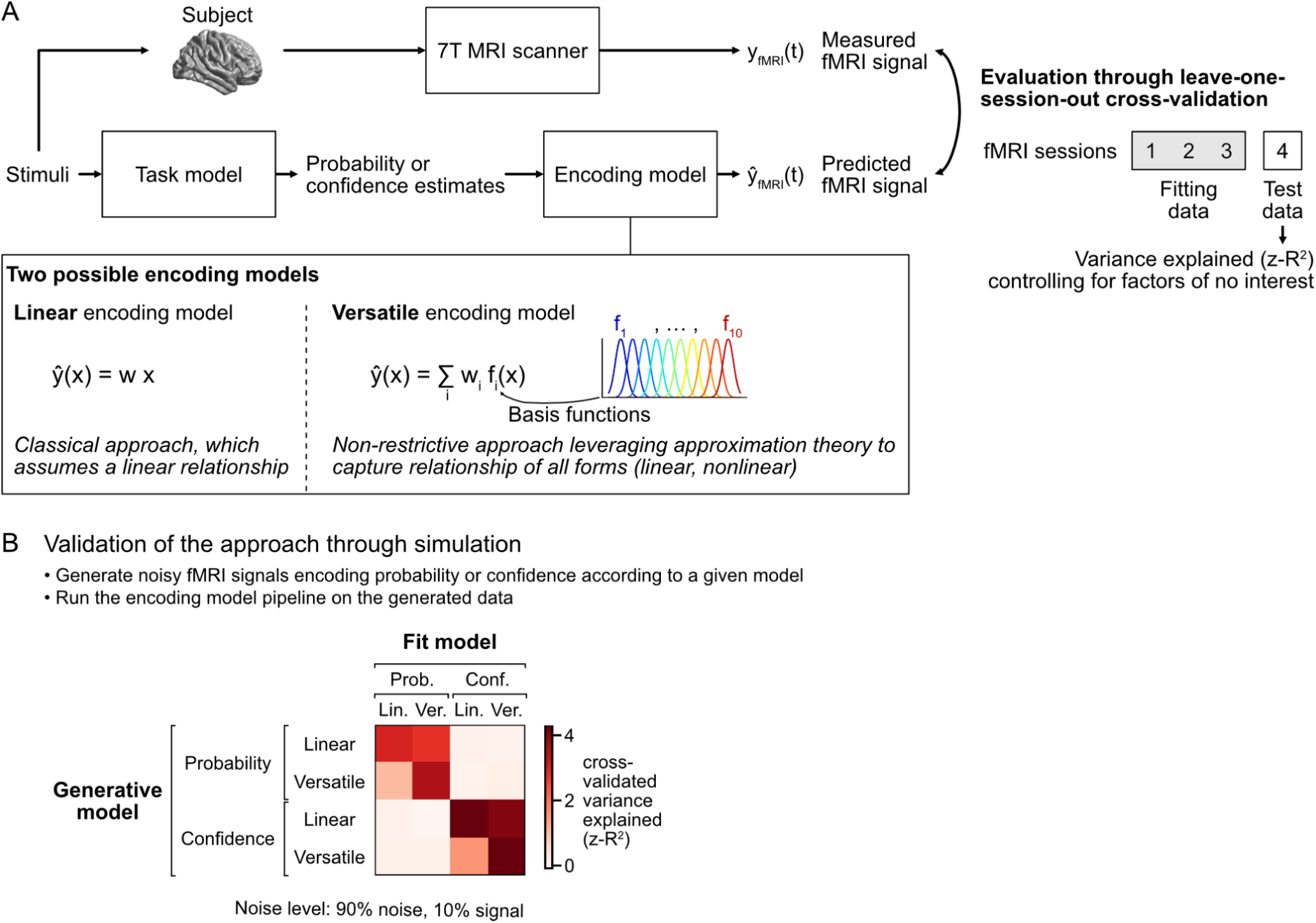
Modeling the encoding of probability or confidence estimates in voxel-wise fMRI signals. (A) Schematic of the encoding models and their evaluation against fMRI data. For each session, the sequence of stimuli presented to the subject is given to the task model to obtain *x*, the probability or confidence estimates on each trial. These estimates are then given to the encoding model to predict *y*, the fMRI signal time series in a voxel. Two classes of encoding models were tested: the linear class, in which the fMRI signal *y* in a given voxel is a linear function of the estimate *x*, and the versatile class, in which the fMRI signal in a given voxel is a weighted sum of basis functions (*f_i_*) that can approximate linear and nonlinear functions of *x*. To evaluate the models, we used a cross-validation procedure in which three out of four sessions were used to fit the encoding model, and the left-out session was used to measure the predictive accuracy of the model (using the coefficient of determination) after controlling for factors of no interest. (B) Simulation results validating the approach end-to-end. Noisy fMRI data for one experiment were generated assuming a given model of neural activity, and the generated data were used to evaluate each of the encoding models using the procedure described in (A). The matrix shows the average score obtained for each evaluated model and each possible generative model across simulated experiments. Linear models explain the data well only when the generative model is linear, whereas versatile models explain the data well for both classes of generative model. Probability and confidence are well separated by the models.

To evaluate the encoding models, we incorporated them into a larger pipeline to obtain predictions at the level of the fMRI signal measured in each voxel (Fig. 2A). The resulting model predicts the fMRI time series in a given session from the probability or confidence estimates, as well as several factors of no-interest that we aim to control for, including: the stimulus onset, the surprise elicited by the stimulus, the entropy associated with the probability estimate (a measure of unpredictability), factors related to the report periods, and motion factors.

We fitted the model parameters and tested the fitted model on independent data using a leave-one-session-out cross-validation procedure (see Methods). During testing, we only kept the part of the model corresponding to the factor of interest (probability or confidence) and calculated the *R^2^* score, which represents the predictive accuracy of the model. Finally, to ensure that a positive score can only be obtained by an encoding related to the sequence specifically observed by the subject and not merely to the statistical structure of the sequences in general, we calculated a null distribution of *R^2^* scores by performing the same analysis after replacing the true sequence with other sequences, and standardized the score obtained with the true sequence relative to this null distribution to yield the final score, which we call *z-R^2^*.

We validated our approach end-to-end using simulations (Fig. 2B). By simulating an experiment following our protocol under the hypothesis that neural activity noisily encodes one of the two types of estimate (probability or confidence) according to an encoding belonging to one of the two classes (linear or versatile), and by applying our fMRI analysis pipeline to the simulated fMRI signal, we established that when neural activity encodes the same type of estimate as the model, the linear model only explains the signal well when this encoding is linear, while the versatile model explains the signal well no matter whether this encoding is linear or not. When neural activity and the model encode a different type of estimate, the model does not explain the signal.

### Neural encoding of probability

The versatile model revealed a significant encoding of probabilities in the prefrontal and parietal cortex (Fig. 3 and Table 1). In contrast, the linear model did not reveal any encoding of probabilities across the cortex, even below the significance threshold (see Fig. 3). The superiority of the versatile encoding model over the linear one is confirmed by the statistical map of the difference in *R^2^* the two models (Supplementary Fig. 1).

**Figure 3.**
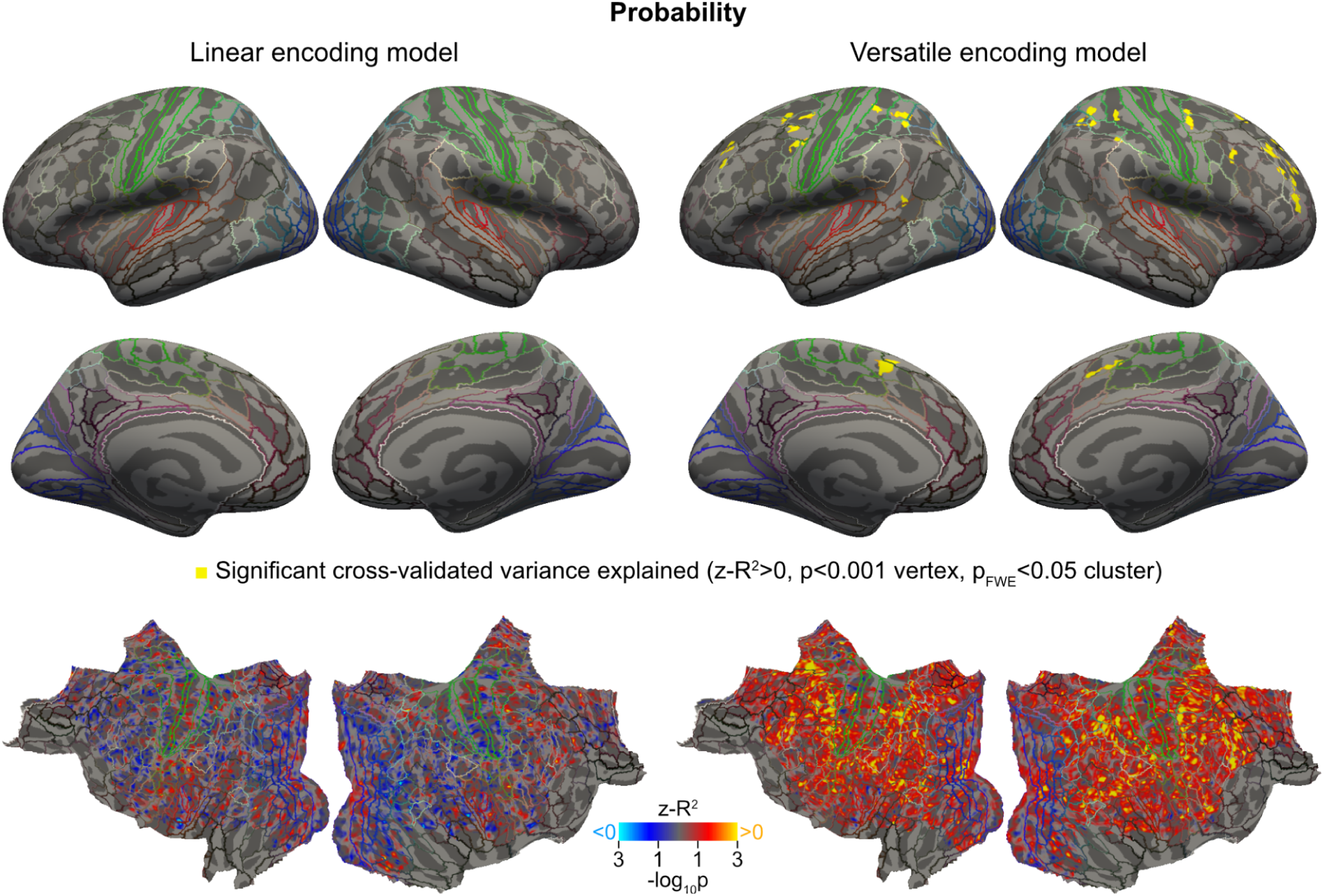
A neural code for probabilities revealed by whole-cortex analysis in prefrontal and parietal cortex, that only the versatile model is able to explain. Cortical maps above show the significance of the cross-validated performance of the linear and versatile encoding models. Top maps show the significant regions after thresholding at p<0.001 and FWE cluster correction at p_FWE_ < 0.05. Bottom maps show the p-values without thresholding or correction on a flattened view of the cortex. P-values correspond to the group-level significance of *z-R^2^* scores obtained across subjects (cold colors for negative scores, hot for positive scores). The HCP-MMP1.0 parcellation is indicated by colored lines (Glasser et al., 2016).

**Table 1.** Significant clusters explained by the versatile encoding model for probability. The names of the cortical areas and parcels refer to the HCP-MMP1.0 atlas (Glasser et al., 2016). Peak x, y, z are MNI coordinates.

| Size (mm<sup>2</sup>) | Num. vertices | Hemisp here | Lobe | Cortical area | Parcel | Peak x | Peak y | Peak z | Peak -log10(p) | Cluster p<sub>FWE</sub> |
| --- | --- | --- | --- | --- | --- | --- | --- | --- | --- | --- |
| 160.4 | 232 | Right | Fr | Dorsolateral_Prefrontal | a9-46v_R | 38.9 | 45.2 | 16 | 6.4097 | 0.0002 |
| 157.4 | 183 | Left | Occ | Early_Visual | V2_L | -25.1 | -97.9 | -2.7 | 4.8187 | 0.0002 |
| 139.01 | 236 | Left | Fr | Paracentral_Lobular_and_Mid_Cingulate | SCEF_L | -9.5 | 5.3 | 55.9 | 4.8964 | 0.0002 |
| 108.66 | 227 | Right | Fr | Premotor | FEF_R | 42.2 | -5.8 | 50.7 | 6.6067 | 0.0002 |
| 106.8 | 274 | Right | Par | Superior_Parietal | LIPv_R | 27.7 | -52.8 | 47.4 | 5.211 | 0.0002 |
| 101.02 | 198 | Left | Par | Superior_Parietal | 7PC_L | -37.4 | -50.2 | 55.6 | 4.5741 | 0.0002 |
| 100.41 | 202 | Right | Fr | Dorsolateral_Prefrontal | 46_R | 27.3 | 29.6 | 36.2 | 4.0095 | 0.0002 |
| 98.83 | 176 | Left | Fr | Premotor | 6a_L | -29.1 | 0.6 | 48.6 | 5.3333 | 0.0002 |
| 80.1 | 169 | Right | Par | Superior_Parietal | 7PC_R | 35.1 | -49.1 | 60.5 | 4.1947 | 0.0006 |
| 71.32 | 161 | Left | Par | Inferior_Parietal | IP1_L | -31.3 | -64.2 | 40.1 | 5.3758 | 0.0014 |
| 70.44 | 127 | Left | Fr | Premotor | FEF_L | -37.3 | -7.6 | 44.5 | 4.6641 | 0.0014 |
| 69.62 | 143 | Right | Fr | Paracentral_Lobular_and_Mid_Cingulate | 24dd_R | 9.5 | 1.1 | 53.9 | 4.8056 | 0.0008 |
| 69.09 | 132 | Left | Fr | Dorsolateral_Prefrontal | 8Ad_L | -25.1 | 24.6 | 33.7 | 5.6881 | 0.0014 |
| 67.68 | 105 | Right | Fr | Dorsolateral_Prefrontal | s6-8_R | 21.7 | 20 | 53.1 | 4.3053 | 0.0008 |
| 64.72 | 136 | Left | Fr | Premotor | PEF_L | -51.2 | -2.6 | 40.4 | 4.6327 | 0.002 |
| 55.09 | 83 | Right | Fr | Dorsolateral_Prefrontal | 9p_R | 19.4 | 38.9 | 37 | 5.5244 | 0.00599 |
| 53.88 | 95 | Right | Fr | Dorsolateral_Prefrontal | 46_R | 35.1 | 33.9 | 27.9 | 4.6328 | 0.00719 |
| 51.51 | 93 | Right | Fr | Dorsolateral_Prefrontal | 9-46d_R | 30.3 | 38.8 | 20.9 | 4.7414 | 0.00918 |
| 49.07 | 57 | Left | Occ | Primary_Visual | V1_L | -17.4 | -100.9 | 0.2 | 4.189 | 0.01296 |
| 48.56 | 70 | Left | Fr | Dorsolateral_Prefrontal | 46_L | -39.2 | 31.5 | 29.3 | 4.2904 | 0.01355 |
| 48.14 | 103 | Left | Par | Somatosensory_and_Motor | 2_L | -36.9 | -35.2 | 56.7 | 4.5609 | 0.01415 |
| 45.65 | 96 | Right | Fr | Dorsolateral_Prefrontal | 8C_R | 33.9 | 10.8 | 33.4 | 4.4846 | 0.0203 |
| 44.86 | 69 | Right | Fr | Dorsolateral_Prefrontal | 9-46d_R | 25.4 | 38.8 | 32.1 | 4.2132 | 0.02247 |
| 43.94 | 88 | Left | Fr | Premotor | 55b_L | -45.1 | 1.5 | 46.8 | 3.7926 | 0.02366 |
| 43.3 | 56 | Right | Fr | Dorsolateral_Prefrontal | 9-46d_R | 25.2 | 45.1 | 29.1 | 4.2352 | 0.02702 |
| 43.14 | 84 | Right | Par | Inferior_Parietal | IP2_R | 50.1 | -37 | 45.4 | 5.2945 | 0.02721 |
| 41.56 | 123 | Right | Par | Somatosensory_and_Motor | 2_R | 31.5 | -35.1 | 46.2 | 3.9812 | 0.03292 |
| 41.06 | 91 | Left | Par | Superior_Parietal | AIP_L | -31.1 | -48.1 | 44.3 | 4.2868 | 0.0345 |
| 38.14 | 72 | Left | Par | Temporo-Parieto-Occipital_Junction | STV_L | -53.8 | -46.6 | 11.7 | 4.2271 | 0.04879 |

This pattern of results is consistent with those predicted under the hypothesis that neural activity encodes probabilities in a nonlinear way (Fig. 2B, second row and first two columns of the matrix). Note that these results do not depend much on the specific choice of basis functions, provided that they have the same approximation properties (see Supplementary Fig. 2).

### Neural encoding of confidence

Contrary to probabilities, for confidence, the linear encoding model significantly explains the measured signal in large regions around the intraparietal and precentral sulci (Fig. 4 and Supplementary Table 1). This is consistent with previous studies on the neural correlates of confidence (Bounmy et al., 2023; Meyniel & Dehaene, 2017). As expected, the versatile encoding model also captures the signal in the regions explained by the linear model. However, unlike probabilities, the regions captured by the versatile and the linear encoding models are very similar throughout the cortex (Fig. 4). This indicates that probability and confidence are encoded differently by the brain, which we sought to further characterize next.

**Figure 4.**
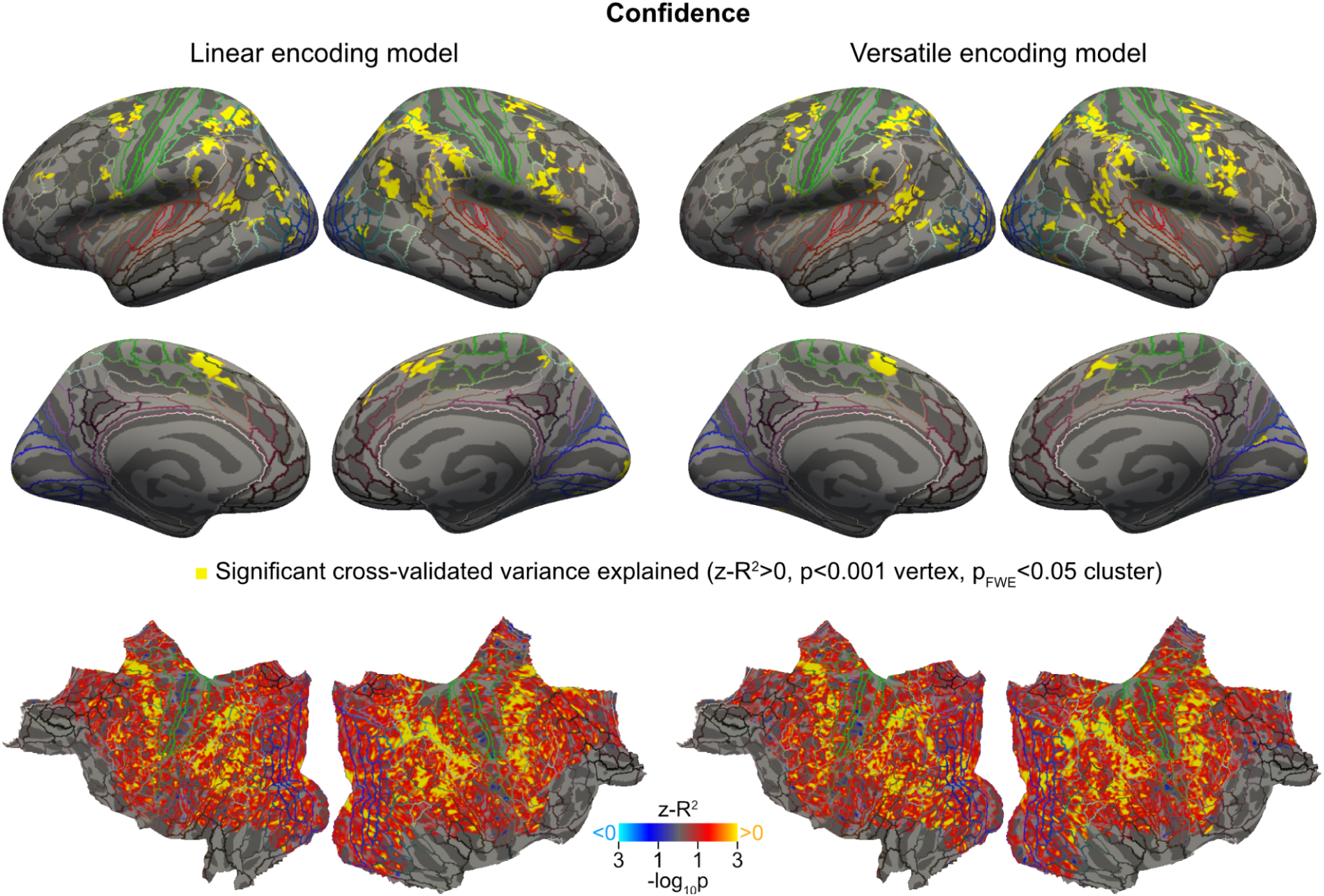
Neural code for confidence. Both the linear and the versatile models explain the neural encoding of confidence in approximately the same regions of the cortex. Plotting conventions are as in Fig. 3.

### Univariate characterization of the neural code

To characterize the neural code, we reconstructed the tuning curves measured at the vertex level. These curves were obtained by calculating the sum of the basis functions weighted by the fitted weights of the versatile model (Fig. 5A). For each subject, we focused on a set of vertices where the predictive accuracy of the versatile model was large enough to ensure a reliable characterization of the tuning curves, which we verified by estimating the tuning curves independently on two halves of the data (Pearson correlation of the estimated weights on the two halves for the vertices of interest: 0.55±0.03 and 0.51±0.04 for probability and confidence, respectively). See examples of tuning curves estimated on the whole and on two halves of the data in Fig. 5B.

**Figure 5.**
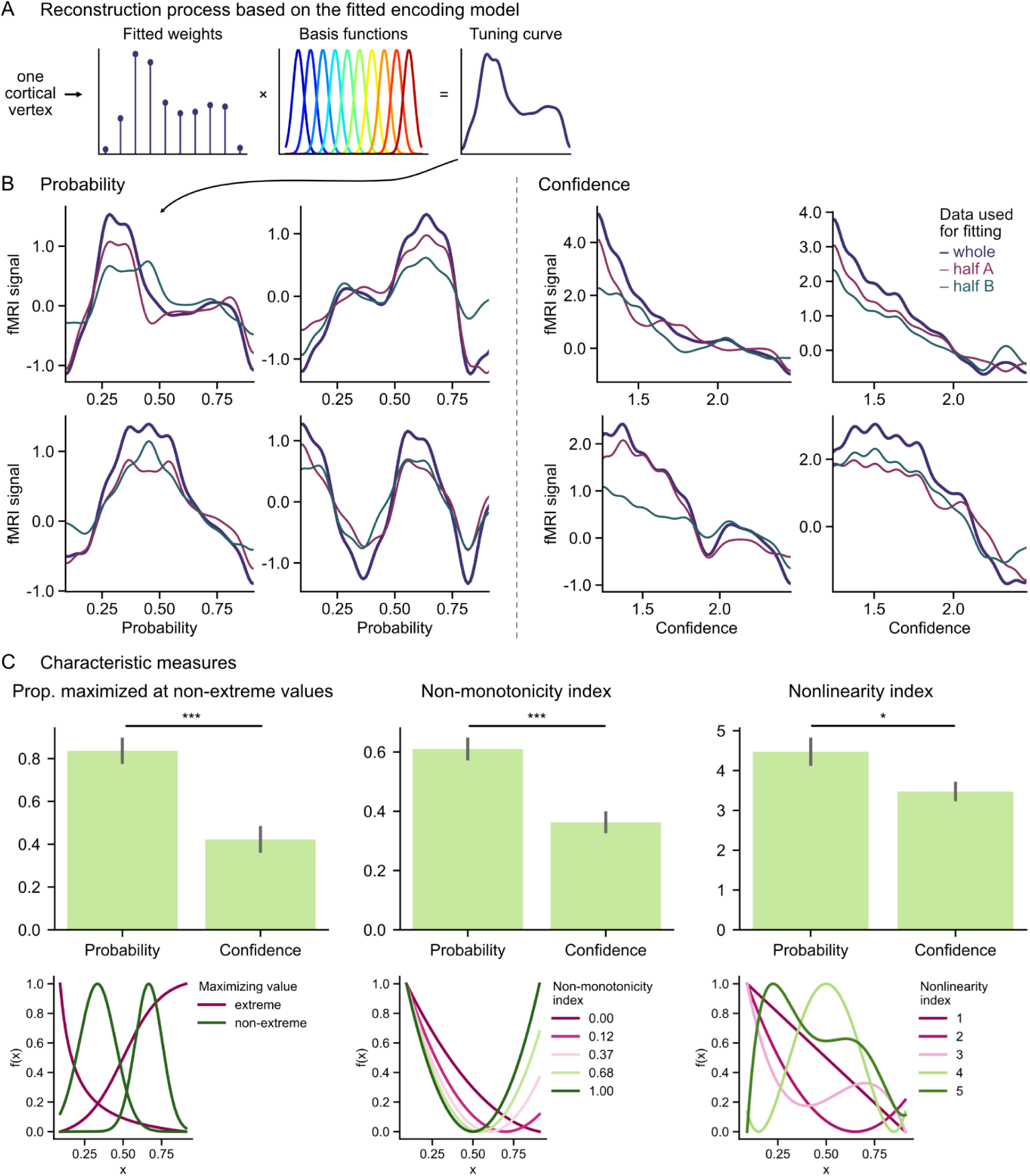
Characterization of the neural encoding of probability and confidence. (A) Schematic of the reconstruction of the tuning curve at the level of a cortical vertex. The weights of the versatile encoding model fitted on the vertex data are used to calculate the tuning curve function, which is equal to the weighted sum of the basis functions. (B) Tuning curves obtained for probability (left) and confidence (right) for example vertices and subjects (out of 30,000 and 120,000 examples, respectively). Within each panel, multiple curves correspond to multiple estimates of the tuning curve for a same vertex: in purple, the curve estimated with the weights fitted on the whole data, red and green, estimated on two independent halves of the data, illustrating test-retest reliability. (C) Quantitative description and comparison of the tuning curves for probability and confidence according to three characteristic measures. The tuning curves for probability are more frequently maximized at non-extreme values, have a higher non-monotonicity index, and a higher nonlinearity index than those for confidence. Bar heights and error bars show mean±s.e.m across subjects. */***: p<0.05/p<0.001, two-tailed tests. Bottom graphs are illustrations of each of the three measures (*x* denotes the encoded variable, which could be probability or confidence, and *f* a tuning curve function).

We used three characteristic measures to describe and compare the tuning curves for probability and confidence (Fig. 5C). For all three measures, the neural encoding of probability and confidence differed significantly. The first measure looked at the location of the maximum of the tuning curve (i.e., the probability or confidence value that maximizes neural activity). This maximum is expected to be close to one of the two extremes in the case of a linear code. Compared to confidence, tuning curves for probability were more often maximized at non-extreme values (Fig. 5C, proportion: 84±6 % for probability vs. 42±6 % for confidence, p<0.001, two-tailed t-test). The second measure looked at the degree of non-monotonicity of the tuning curves, quantified by a continuous index between 0 and 1 (see illustration in Fig. 5C). According to this index, tuning curves were more non-monotonic for probability than for confidence (Fig. 5C, 0.61±0.04 for probability vs. 0.36±0.04 for confidence, p<0.001, two-tailed t-test). The third measure looked at the degree of nonlinearity of the tuning curve (corresponding to the degree of a polynomial, the higher the degree, the more nonlinearities). This measure showed that tuning curves were more nonlinear for probability than for confidence (Fig. 5C, 4.5±0.4 for probability vs. 3.5±0.2 for confidence, p<0.05, two-tailed Kruskal-Wallis test by ranks).

### Multivariate characterization of the neural code

So far, the analysis has been univariate, focusing on one voxel at a time. If different voxels are tuned to different ranges of probability, as indicated by the above encoding analysis, then patterns of voxel responses should be informative about probability, which we tested with a multivariate decoding approach. Furthermore, the pattern of voxel response should exhibit different geometric properties if they arise from nonlinear vs. linear codes, providing another test of the hypothesis that the neural code for probability is highly nonlinear. Below, we present thise two analyses.

We measured the extent to which probability (and for comparison, confidence) could be decoded in different brain regions by dividing the cortex following the parcellation by Glasser et al (Glasser et al., 2016) and pooling homologue regions in both hemispheres, resulting in 180 regions. We based our decoding approach on the versatile encoding model presented above. The versatile encoding model quantifies in each voxel the weights of basis functions of probability. The basis functions being equally spaced narrow Gaussian functions, the set of estimated weights can be thought of as characterizing the voxel responses to different bins (i.e. narrow ranges) of probability. We tested whether it is possible to identify (i.e. decode) these probability bins given the pattern of voxel responses that they elicit. We used 5 probability bins instead of 10 as in the encoding approach presented above to make the estimates of weights more reliable, and thus decoding easier. We adopted the same approach to decode confidence levels.

We trained and tested the decoder on different data sets, using leave-one-session-out cross validation at the subject-level (Varoquaux et al., 2017). First, we reduced dimensionality in each region by selecting the 100 voxels that are the most informative about probabilities (using the *z-R²* metric introduced above, but estimated using the training sessions only). Then we estimated the patterns of voxel responses for different probability bins, for the test session on the one hand and the three training sessions together on the other hand. Last, we decoded the probability bin corresponding to a response pattern in the test session by identifying the probability bin eliciting the most similar response pattern in the training session. We computed the decoding accuracy for each brain region and tested for statistical significance against chance-level accuracy at the group level (see Fig. 5 and Methods).

Decoding accuracy was generally larger for confidence than probability (Fig 6, Tables 2 & 3), which is expected given that the versatile encoding model accounted for voxel responses in more regions based on confidence than on probability (Fig. 4 vs. 5). For probability, we found 6 regions with FDR-significant decoding accuracy. They comprised the dorsolateral prefrontal cortex and the intraparietal cortex, which had been identified with the encoding approach. For confidence, more regions exhibited a FDR-significant decoding accuracy, notably in parietal and prefrontal cortex, similar to the regions identified with the encoding approach.

**Figure 6.**
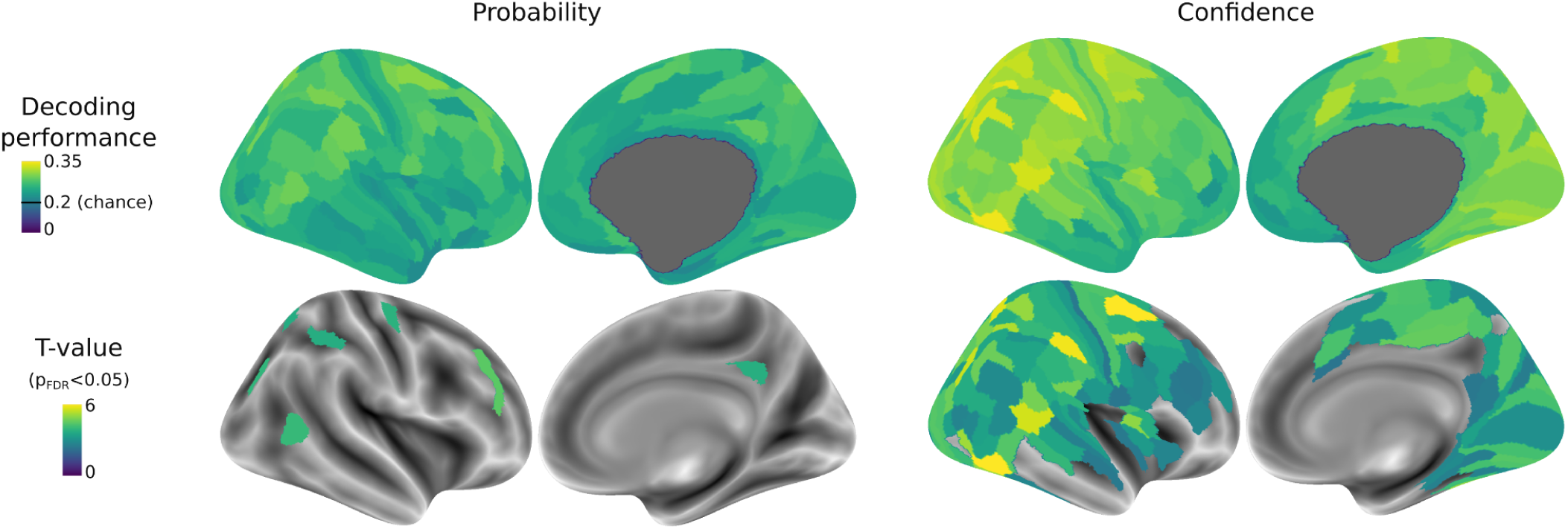
Decoding probability and confidence from the voxel response patterns obtained with the versatile encoding model. Group-level decoding accuracy for each cortical region (180 bihemispheric parcels from the HCP-MMP1.0 atlas, rendered on the right hemisphere for illustration purpose). Top: Mean accuracy across subjects (chance level is one out of five bins, i.e. 0.2). Bottom: T-values of a two-tailed t-test for accuracy different from chance level. Only regions statistically significant after FDR-correction (p<0.05) for multiple comparisons across the 180 regions are displayed (also see Table 2 and 3). The decoding method is based on the similarity (correlation distance) of voxel response patterns estimated by the encoding model in a training set and a test set (see Methods).

**Table 2.** Decoding accuracy for probability. . Names of cortical areas are as in Table 1. The decoding accuracy is compared to chance level (0.2) with a two-sided t-test. Only parcels with significant decoding accuracy (p_FDR_<0.05) are reported.

| Lobe | Cortical area | Parcel | Mean | T value | P value | $p_{\text{FDR}}$ |
| --- | --- | --- | --- | --- | --- | --- |
| Fr | Dorsolateral_Prefrontal | 46 | 0.27 | 4.46 | 0.0002 | 0.0327 |
| Par | Temporo-Parieto-Occipital_Junction | TPOJ2 | 0.27 | 4.03 | 0.0006 | 0.0327 |
| Occ | Inferior_Parietal | IP0 | 0.27 | 3.93 | 0.0007 | 0.0327 |
| Fr | Premotor | 6d | 0.28 | 3.84 | 0.0009 | 0.0327 |
| Par | Posterior_Cingulate | 31pv | 0.27 | 3.76 | 0.0011 | 0.0327 |
| Par | Superior_Parietal | AIP | 0.28 | 3.74 | 0.0012 | 0.0327 |
| Par | Superior_Parietal | VIP | 0.28 | 3.70 | 0.0013 | 0.0327 |

**Table 3.** Decoding accuracy for confidence. . Names of cortical areas are as in Table 1. The decoding accuracy is compared to chance level (0.2) with a two-sided t-test. Only parcels with significant decoding accuracy (p_FDR_<0.05) are reported.

| Lobe | Cortical area | Parcel | Mean | T value | P value | $p_{FDR}$ |
| --- | --- | --- | --- | --- | --- | --- |
| Fr | Premotor | 6a | 0.30 | 6.29 | 0.0000 | 0.0002 |
| Temp | MT+ Complex and Neighboring Visual Areas | PH | 0.34 | 6.18 | 0.0000 | 0.0002 |
| Par | Inferior Parietal | PFt | 0.34 | 5.87 | 0.0000 | 0.0003 |
| Par | Superior Parietal | VIP | 0.32 | 5.72 | 0.0000 | 0.0003 |
| Temp | Temporo-Parieto-Occipital Junction | TPOJ1 | 0.33 | 5.59 | 0.0000 | 0.0004 |
| Occ | Inferior Parietal | IP0 | 0.32 | 5.52 | 0.0000 | 0.0004 |
| Par | Superior Parietal | LIPd | 0.34 | 5.28 | 0.0000 | 0.0006 |
| Occ | MT+ Complex and Neighboring Visual Areas | V4t | 0.31 | 5.15 | 0.0000 | 0.0007 |
| Par | Superior Parietal | 7PC | 0.32 | 5.06 | 0.0000 | 0.0007 |
| Temp | Ventral Stream Visual | FFC | 0.31 | 5.04 | 0.0000 | 0.0007 |
| Fr | Premotor | 55b | 0.31 | 5.02 | 0.0000 | 0.0007 |
| Ins | Insular and Frontal Opercular | FOP3 | 0.29 | 4.99 | 0.0000 | 0.0007 |
| Par | Superior Parietal | MIP | 0.32 | 4.81 | 0.0001 | 0.0010 |
| Par | Temporo-Parieto-Occipital Junction | STV | 0.31 | 4.75 | 0.0001 | 0.0010 |
| Occ | Dorsal Stream Visual | V7 | 0.30 | 4.75 | 0.0001 | 0.0010 |
| Par | Early Auditory | PFcm | 0.30 | 4.75 | 0.0001 | 0.0010 |
| Occ | MT+ Complex and Neighboring Visual Areas | V3CD | 0.30 | 4.66 | 0.0001 | 0.0012 |
| Par | Inferior Parietal | PGs | 0.33 | 4.64 | 0.0001 | 0.0012 |
| Par | Paracentral Lobular and Mid Cingulate | 23c | 0.28 | 4.59 | 0.0001 | 0.0013 |
| Par | Paracentral Lobular and Mid Cingulate | 5mv | 0.28 | 4.52 | 0.0002 | 0.0014 |
| Par | Dorsal Stream Visual | IPS1 | 0.33 | 4.52 | 0.0002 | 0.0014 |
| Occ | Early Visual | V3 | 0.31 | 4.42 | 0.0002 | 0.0016 |
| Occ | Posterior Cingulate | POS2 | 0.29 | 4.40 | 0.0002 | 0.0016 |
| Temp | Lateral Temporal | PHT | 0.30 | 4.40 | 0.0002 | 0.0016 |
| Par | Posterior Cingulate | PCV | 0.30 | 4.35 | 0.0002 | 0.0018 |
| Fr | Paracentral Lobular and Mid Cingulate | 6mp | 0.31 | 4.34 | 0.0003 | 0.0018 |
| Occ | Early Visual | V4 | 0.30 | 4.32 | 0.0003 | 0.0018 |
| Fr | Somatosensory and Motor | 4 | 0.28 | 4.31 | 0.0003 | 0.0018 |
| Fr | Anterior Cingulate and Medial Prefrontal | p32pr | 0.31 | 4.29 | 0.0003 | 0.0018 |
| Par | Inferior Parietal | PFop | 0.29 | 4.28 | 0.0003 | 0.0018 |
| Fr | Paracentral Lobular and Mid Cingulate | 24dd | 0.28 | 4.26 | 0.0003 | 0.0018 |
| Fr | Premotor | 6d | 0.32 | 4.21 | 0.0004 | 0.0019 |
| Occ | Ventral Stream Visual | VMV2 | 0.28 | 4.21 | 0.0004 | 0.0019 |
| Occ | Early Visual | V2 | 0.30 | 4.20 | 0.0004 | 0.0019 |
| Fr | Posterior Opercular | FOP1 | 0.27 | 4.18 | 0.0004 | 0.0020 |
| Temp | Ventral Stream Visual | VVC | 0.30 | 4.12 | 0.0004 | 0.0022 |
| Occ | Ventral Stream Visual | PIT | 0.29 | 4.07 | 0.0005 | 0.0024 |
| Par | Superior Parietal | AIP | 0.30 | 4.00 | 0.0006 | 0.0028 |
| Temp | Auditory Association | STSvp | 0.28 | 3.96 | 0.0007 | 0.0031 |
| Par | Inferior Parietal | PF | 0.29 | 3.89 | 0.0008 | 0.0035 |
| Fr | Premotor | FEF | 0.30 | 3.89 | 0.0008 | 0.0035 |
| Par | Early Auditory | RI | 0.30 | 3.88 | 0.0008 | 0.0035 |
| Par | Somatosensory and Motor | 2 | 0.28 | 3.83 | 0.0009 | 0.0038 |
| Fr | Paracentral Lobular and Mid Cingulate | 6ma | 0.28 | 3.78 | 0.0011 | 0.0043 |
| Par | Paracentral Lobular and Mid Cingulate | 5m | 0.29 | 3.77 | 0.0011 | 0.0043 |
| Fr | Dorsolateral Prefrontal | 8Av | 0.27 | 3.75 | 0.0011 | 0.0044 |
| Fr | Insular and Frontal Opercular | FOP4 | 0.30 | 3.74 | 0.0012 | 0.0044 |
| Par | Temporo-Parieto-Occipital Junction | PSL | 0.30 | 3.73 | 0.0012 | 0.0045 |
| Fr | Dorsolateral Prefrontal | i6-8 | 0.27 | 3.70 | 0.0013 | 0.0047 |
| Occ | Inferior Parietal | PGp | 0.28 | 3.69 | 0.0013 | 0.0048 |
| Par | Superior Parietal | LIPv | 0.30 | 3.67 | 0.0014 | 0.0048 |
| Occ | Temporo-Parieto-Occipital Junction | TPOJ3 | 0.29 | 3.67 | 0.0014 | 0.0048 |
| Par | Superior Parietal | 7AL | 0.31 | 3.66 | 0.0014 | 0.0049 |
| Par | Somatosensory and Motor | 1 | 0.29 | 3.64 | 0.0015 | 0.0050 |
| Par | Superior Parietal | 7PI | 0.29 | 3.62 | 0.0016 | 0.0051 |
| Par | Inferior Parietal | IP1 | 0.29 | 3.55 | 0.0018 | 0.0058 |
| Occ | Dorsal_Stream_Visual | V3A | 0.28 | 3.55 | 0.0018 | 0.0058 |
| Temp | Auditory_Association | STSdp | 0.29 | 3.53 | 0.0020 | 0.0060 |
| Occ | MT+_Complex_and_Neighboring_Visual_Areas | FST | 0.29 | 3.51 | 0.0020 | 0.0060 |
| Par | Inferior_Parietal | PFm | 0.27 | 3.51 | 0.0020 | 0.0060 |
| Temp | Early_Auditory | PBelt | 0.28 | 3.51 | 0.0020 | 0.0060 |
| Par | Posterior_Cingulate | 7m | 0.29 | 3.44 | 0.0024 | 0.0070 |
| Occ | MT+_Complex_and_Neighboring_Visual_Areas | MST | 0.27 | 3.44 | 0.0024 | 0.0070 |
| Par | Posterior_Cingulate | ProS | 0.27 | 3.42 | 0.0026 | 0.0072 |
| Fr | Dorsolateral_Prefrontal | 8C | 0.27 | 3.41 | 0.0026 | 0.0073 |
| Occ | MT+_Complex_and_Neighboring_Visual_Areas | LO1 | 0.28 | 3.39 | 0.0027 | 0.0074 |
| Fr | Inferior_Frontal | IFJa | 0.27 | 3.39 | 0.0027 | 0.0074 |
| Temp | Early_Auditory | A1 | 0.27 | 3.35 | 0.0030 | 0.0079 |
| Fr | Premotor | 6v | 0.27 | 3.32 | 0.0032 | 0.0084 |
| Occ | Posterior_Cingulate | DVT | 0.30 | 3.32 | 0.0033 | 0.0084 |
| Temp | Auditory_Association | A4 | 0.27 | 3.25 | 0.0038 | 0.0097 |
| Ins | Insular_and_Frontal_Opercular | MI | 0.26 | 3.25 | 0.0039 | 0.0097 |
| Par | Somatosensory_and_Motor | 3b | 0.28 | 3.21 | 0.0042 | 0.0103 |
| Occ | Dorsal_Stream_Visual | V3B | 0.27 | 3.20 | 0.0043 | 0.0106 |
| Par | Posterior_Opercular | OP4 | 0.27 | 3.19 | 0.0044 | 0.0107 |
| Par | Inferior_Parietal | IP2 | 0.26 | 3.13 | 0.0051 | 0.0121 |
| Fr | Dorsolateral_Prefrontal | 9-46d | 0.27 | 3.08 | 0.0058 | 0.0136 |
| Fr | Paracentral_Lobular_and_Mid_Cingulate | SCEF | 0.28 | 3.05 | 0.0061 | 0.0141 |
| Fr | Premotor | 6r | 0.27 | 3.01 | 0.0068 | 0.0155 |
| Par | Paracentral_Lobular_and_Mid_Cingulate | 5L | 0.27 | 2.99 | 0.0072 | 0.0160 |
| Occ | Primary_Visual | V1 | 0.28 | 2.98 | 0.0072 | 0.0160 |
| Par | Superior_Parietal | 7Am | 0.27 | 2.95 | 0.0078 | 0.0169 |
| Fr | Paracentral_Lobular_and_Mid_Cingulate | 24dv | 0.26 | 2.95 | 0.0078 | 0.0169 |
| Occ | Ventral_Stream_Visual | V8 | 0.27 | 2.93 | 0.0081 | 0.0173 |
| Fr | Inferior_Frontal | IFJp | 0.28 | 2.93 | 0.0082 | 0.0174 |
| Par | Temporo-Parieto-Occipital_Junction | TPOJ2 | 0.28 | 2.90 | 0.0088 | 0.0185 |
| Temp | Medial_Temporal | PHA1 | 0.27 | 2.87 | 0.0093 | 0.0193 |
| Par | Inferior_Parietal | PGi | 0.27 | 2.86 | 0.0096 | 0.0196 |
| Temp | Early_Auditory | LBelt | 0.27 | 2.85 | 0.0098 | 0.0198 |
| Occ | MT+_Complex_and_Neighboring_Visual_Areas | MT | 0.28 | 2.81 | 0.0107 | 0.0215 |
| Par | Posterior_Opercular | OP1 | 0.25 | 2.79 | 0.0114 | 0.0223 |
| Temp | Medial_Temporal | PreS | 0.27 | 2.78 | 0.0115 | 0.0223 |
| Fr | Inferior_Frontal | IFSa | 0.27 | 2.78 | 0.0115 | 0.0223 |
| Temp | Lateral_Temporal | TE2p | 0.26 | 2.75 | 0.0124 | 0.0237 |
| Fr | Insular_and_Frontal_Opercular | AVI | 0.26 | 2.74 | 0.0127 | 0.0240 |
| Occ | MT+_Complex_and_Neighboring_Visual_Areas | LO3 | 0.28 | 2.71 | 0.0135 | 0.0254 |
| Occ | Dorsal_Stream_Visual | V6A | 0.27 | 2.69 | 0.0142 | 0.0264 |
| Temp | Medial_Temporal | PHA2 | 0.27 | 2.62 | 0.0164 | 0.0301 |
| Occ | Ventral_Stream_Visual | VMV3 | 0.26 | 2.61 | 0.0168 | 0.0301 |
| Fr | Dorsolateral_Prefrontal | 46 | 0.26 | 2.61 | 0.0168 | 0.0301 |
| Temp | Auditory_Association | A5 | 0.27 | 2.61 | 0.0169 | 0.0301 |
| Occ | Ventral_Stream_Visual | VMV1 | 0.27 | 2.56 | 0.0190 | 0.0335 |
| Occ | Dorsal_Stream_Visual | V6 | 0.26 | 2.53 | 0.0202 | 0.0352 |
| Par | Posterior_Cingulate | POS1 | 0.26 | 2.51 | 0.0210 | 0.0363 |
| Temp | Auditory_Association | STSda | 0.26 | 2.46 | 0.0237 | 0.0405 |
| Fr | Inferior_Frontal | IFSp | 0.25 | 2.45 | 0.0238 | 0.0405 |
| Ins | Insular_and_Frontal_Opercular | Pol2 | 0.26 | 2.45 | 0.0241 | 0.0405 |
| Fr | Somatosensory_and_Motor | 3a | 0.25 | 2.43 | 0.0250 | 0.0417 |
| Par | Posterior_Cingulate | v23ab | 0.26 | 2.40 | 0.0266 | 0.0439 |
| Fr | Dorsolateral_Prefrontal | p9-46v | 0.25 | 2.40 | 0.0269 | 0.0441 |
| Fr | Posterior_Opercular | 43 | 0.25 | 2.37 | 0.0284 | 0.0461 |
| Fr | Anterior_Cingulate_and_Medial_Prefrontal | a24pr | 0.26 | 2.34 | 0.0304 | 0.0489 |

Decoding accuracy indicates that patterns of voxel activity are informative about probability or confidence, but it does not characterize the type of neural code being used. We examined more closely the patterns of voxel responses to test for the existence of different codes for probability and confidence. More precisely, we examined the matrices that quantify the dissimilarity of patterns of voxel responses between bins of probability (and similarly, bins of confidence) that served as a basis for decoding. These matrices are called representational dissimilarity matrices (RDM) (Kriegeskorte & Kievit, 2013). Different types of code predict different types of RDM. If the code is highly nonlinear and non-monotonic, as suggested by the encoding analysis of probability, then the patterns of voxel responses should be maximally similar when representing the same probability bin, and equally dissimilar between a given bin and any other bin, resulting in an identity RDM (Fig. 7A). In contrast, if the code is highly linear (or more generally monotonic), as suggested by the encoding analysis of confidence, then the patterns of voxel responses should be more similar for confidence bins that are closer to one another, resulting in a graded RDM. To determine which code best accounted for the empirical RDMs, we regressed the RDMs for probability (and confidence) obtained in each region onto the identity RDM and graded RDM arising from highly nonlinear and linear codes, respectively.

**Figure 7:**
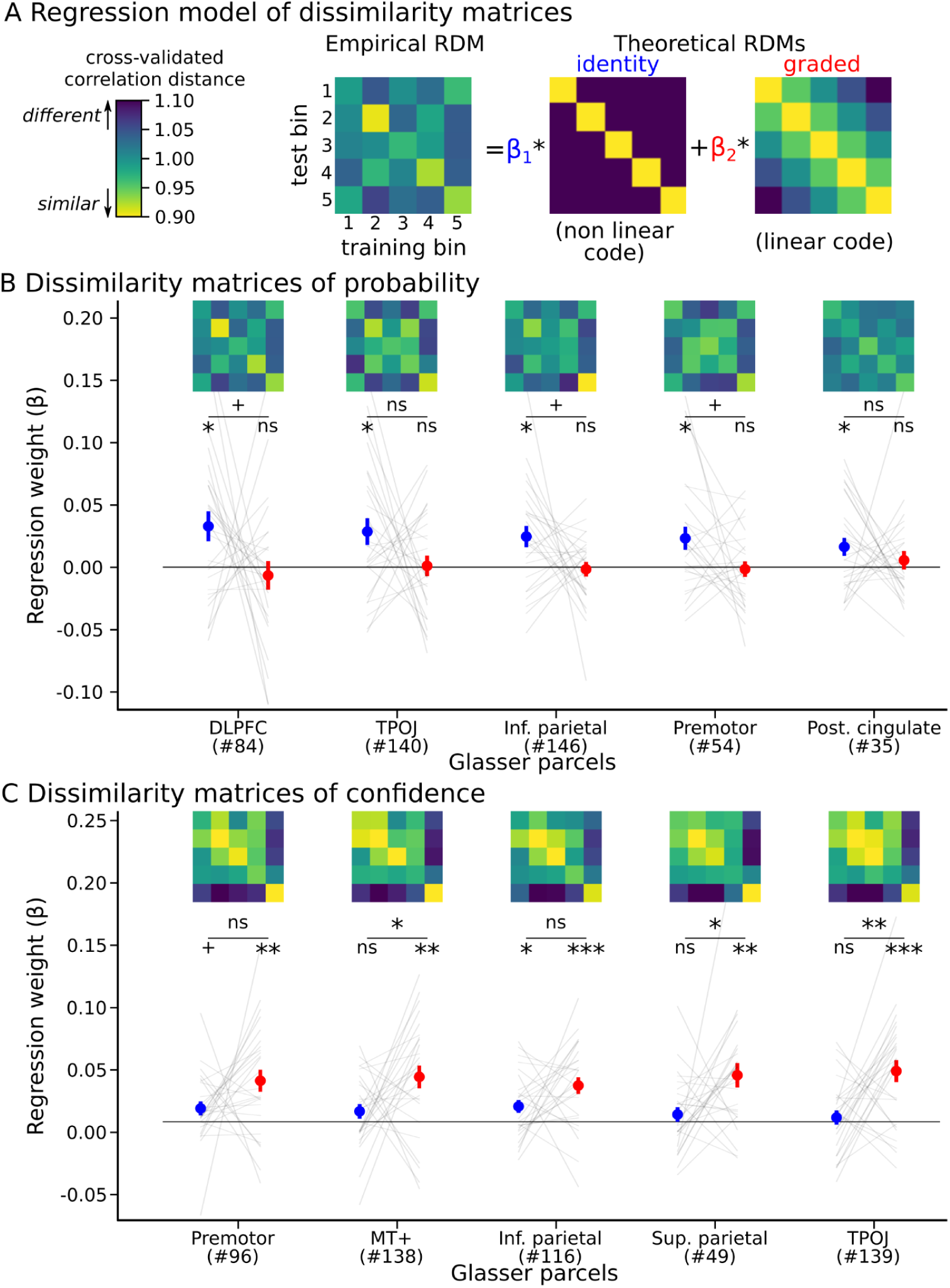
The dissimilarity of voxel response patterns across bins of probability and confidence supports the existence of different types of codes. (A) The representational dissimilarity matrices (RDM) that served for the decoding analysis (Fig. 6) were averaged across test sessions and analyzed with a regression analysis at the subject-level. Two theoretical RDMs were considered in the regression: the “identity” RDM, that should result from a highly nonlinear (and non-monotonic) code, and the “graded” RDM, that should result from a linear (or more generally, monotonic) code. (B) Average RDM of probability levels and regression results in each of the 5 regions with top decoding significance for probability. Each line corresponds to a participant; colored dots and error bars show the average and S.E.M. (C) Same for confidence. In B and C, the same color bar (shown in A) is used for all RDMs. Each set of regression coefficients was tested at the group level against 0 (two-tailed t-test) and compared with a paired t-test (two-tailed); ns: p>0.1; +: p<0.1; *: p<0.05; **:p<0.005; ***:p<0.0005. The parcel numbers correspond to the Glasser parcellation (Glasser et al., 2016); 84: Dorsolateral Prefrontal, Area 46; 140: Temporo-Parieto-Occipital Junction, Area 2, 146: Inferior Parietal, Area 0; 54: Premotor, Dorsal Area 6d, 35: Posterior Cingulate, area 31pv; 96: Premotor, Area 6 anterior; 138: MT+ Complex and Neighboring Visual Areas, Area PH; 116: Inferior Parietal, Area PFt; 49: Superior Parietal, Ventral IntraParietal Complex, 139: Temporo-Parieto-Occipital Junction, Area 1.

We first focused on the 5 regions exhibiting the most significant decoding accuracy, separately for probability (Fig. 7B) and confidence (Fig. 7C). The identity RDM significantly accounted for the RDMs for probability, while no significant effect of the graded RDM was found, and the difference between the two approached significance in most regions. In contrast, for confidence, the graded RDM significantly accounted for the RDMs, the identity RDM yielded no significant effect, and their difference was significant in most regions.

We then carried another analysis that covered all regions. We counted the number of regions best explained by the identity or the graded RDM based on the maximum regression coefficient. A majority of regions followed the graded RDM in the case of confidence (M_graded_=0.672, SEM=0.132, t=3.6, p=0.0014, t-test against 0.5) whereas a majority of regions followed the identity RDM in the case of probability (M_graded_=0.436, SEM=0.086, t=-1.8, p=0.078, t-test against 0.5), and the difference between confidence and probability was significant (paired difference 0.236, SEM=0.0462, t=3.6, p=0.0014, t-test). Together, these results indicate that the codes for probability and confidence are different, being respectively highly nonlinear and mostly linear, which confirms the conclusion of the encoding analysis presented above.

## Discussion

We used a task in which participants estimated the latent probability of a stimulus occurring in a sequence. Subjects accurately tracked this latent probability, as revealed by the comparison of their reports with an ideal observer. We identified a representation of this latent probability in the fMRI signals recorded outside of the report periods, particularly in the dorsolateral prefrontal cortex and intraparietal cortex. Crucially, this representation was based on a nonlinear code. In contrast, the representation of the confidence that accompanied the probability estimate was based on a linear code. Detailed analysis revealed that the vast majority of fMRI voxel tuning curves for probability were non-monotonic, with one or more local extrema. In contrast, the tuning curves for confidence were essentially linear. In addition, a linear code and a highly nonlinear code are expected to result in different geometries, which we confirmed with a multivariate analysis of the patterns of voxels responses to probability and confidence.

Our results relied on the use of a versatile encoding model capable of accommodating tuning curves of almost any shape. To this end, we combined the universal function approximation properties of basis sets (Bishop, 2007; Franke, 1982) with the use of linearizing encoding models for fMRI (Huth et al., 2016; Naselaris et al., 2011). This method consists of applying basis functions to a quantity of interest to obtain features and then modelling the fMRI signal in each voxel as a linear combination of these features, following the general linear model approach that is massively used in fMRI (Friston et al., 2007). This approach is related to other methods that can accommodate nonlinear tuning curves, such as population receptive field (pRF) mapping (Barretto-García et al., 2023; Dumoulin & Wandell, 2008; Harvey et al., 2013). The key difference is that pRF methods assume a specific form of nonlinearity, typically bell-shaped tuning curves, corresponding to the idea that a voxel is selective for a range of values. In contrast, our method can accommodate tuning curves exhibiting a selectivity for multiple value ranges. It turns out that approximately a third of tuning curves we obtained for probability did not conform to a bell-shaped tuning as they exhibited more than one peak. Our finding that the neural representation of probability is highly nonlinear retrospectively explains null findings in previous studies that used methods assuming a linear code (Bounmy et al., 2023; Lebreton et al., 2015; Marshall et al., 2022).

Our results raise the puzzling question of why some quantities are encoded with linear codes, such as confidence here, or reward (Lebreton et al., 2009; Padoa-Schioppa & Assad, 2008), salience (Kutlu et al., 2021), surprise (Meyniel & Dehaene, 2017; O’Reilly et al., 2013), prediction error (Pessiglione et al., 2006; Schultz et al., 1997), evidence accumulation (Brunton et al., 2013; Gold & Shadlen, 2007) and some other quantities are encoded with nonlinear codes, such as probability here, or orientation of visual object (Hubel & Wiesel, 1959), angle in arm reaching (Georgopoulos et al., 1986), numerosity (Nieder, 2016). This question is beyond the scope of our study, but we mention some speculations. Some scalar quantities lie on an axis whose direction is relevant to the regulation of behavior and brain processes. For instance, humans and other animals generally seek more, not less, rewards (Rangel et al., 2008). A linear code with increasing activity levels for larger rewards may, in downstream circuits, facilitate the invigoration of behavior to obtain larger rewards (Pessiglione et al., 2007), and the comparison of different reward levels (which, in a linear code, simply amounts to comparing activity levels). A similar argument applies to salience, surprise, accumulated evidence, and lack of confidence. Higher values of these quantities usually enhance other processing: more salient and surprising events elicit stronger orienting responses (Sara & Bouret, 2012), lower confidence about a learned estimate increases the learning rate (Foucault & Meyniel, 2023; Nassar et al., 2010). In contrast, in our task, the probability of occurrence of a right- vs. left-tilted Gabor patch has no valence and it is not immediately relevant to behavior. Interestingly, the average response across a population (of neurons or voxels) is invariant to the value encoded in a nonlinear code with sufficiently diverse tuning curves. Assuming that more neural activity is costly (Gallistel, 2017), this invariance property implies that the same energy budget is expended to represent any probability, in particular for low (close to 0) and large (close to 1) probabilities. In contrast, encoding reward, salience, surprise, or lack of confidence with increasing (linear) levels of activity appropriately expends more energy on larger, behaviorally relevant values.

Here, we found a neural representation of probabilities predominantly in the dorsolateral prefrontal cortex, the precentral sulcus and the parietal region. The dorsolateral prefrontal and intraparietal cortices have been reported to host a general coding system for magnitudes of different types, from number of objects to proportions in humans and monkeys (Eger, 2016; Nieder, 2016). Our results suggest that this general coding system may also encode probability. In our results, the dorsolateral portion of the prefrontal cortex appeared to encode only probability, whereas the precentral sulcus and the parietal regions encoded both confidence (with a linear code) and probability (with a nonlinear code). Simulations showed that a linear code for confidence and a nonlinear code for probability can be clearly distinguished from one another; their co-localization is therefore a notable finding. We speculate that if a region is involved in estimating and encoding of probabilities, it should be more active when the estimate is updated more, which typically occurs when confidence is lower (Bounmy et al., 2023; McGuire et al., 2014; Tomov et al., 2020). Co-localization of a nonlinear code for probability and a decreasing (linear) code for confidence would then be expected. Future studies should distinguish between updating and representing probabilities, and our results suggest that these processes may differentially involve the dorsolateral prefrontal cortex on the one hand, and the precentral sulcus and parietal region on the other hand.

It is important to distinguish between the representation of an event probability and the representation of a probability density function (of scalar variables such as the orientation of a grating). The latter has been investigated by two lines of research focusing on encoding and decoding, respectively. On the one hand, some researchers have proposed that probability density functions are encoded in activity patterns (Jazayeri & Movshon, 2006; Zemel et al., 1998). On the other hand, research on probabilistic population codes (Ma et al., 2006) posits that, with certain models of neural variability, probability density functions can be decoded from activity patterns in the form of a posterior distribution over some scalar variable. Here, we have identified a neural code for the event probability (which is the mean of the posterior distribution); it remains open whether the brain also codes a probability density function of the event probability (i.e. its full posterior distribution, see Eq. 1 in Methods). Following the encoding approach (à la Zemel et al), we have tested a variant of the versatile encoding model that encodes the full posterior distribution of the event probability (instead of its mean; see Methods). We found no clear evidence for such a representation (see Supplementary Fig. 3). We did not attempt to follow the probabilistic population code approach (à la Ma et al), because it requires modelling the variability of the responses elicited by a given probability (van Bergen et al., 2015), which is difficult to obtain in practice but promising for future research.

We acknowledge that our results do not exclude the possibility that other coding schemes are used to represent probability. Functional MRI can study rate codes (in which information is conveyed by the rate of spikes, not their timing) at the level of voxels, but its temporal resolution is incompatible with the study of temporal codes (Dayan & Abbott, 2005), especially at the level of neurons. Temporal codes may also be used for probabilities. Probabilities could in principle also be stored in synaptic weights (Iigaya, 2016) or intracellular substrates (Gallistel, 2017). It has been claimed that the function of neural activity (and thus indirectly, the fMRI signal (Logothetis et al., 2001)) is to transmit information, which is much more energetically costly than storing information in a cellular substrate (Gallistel, 2017). We note that the probability here has to be constantly updated because there are change points occurring at random, un-signalled times. The reason why we found a representation of probabilities in fMRI activity patterns may be because probabilities were being constantly updated by subjects. This updating process, which involves the interaction of neurons that combine the current estimate with information about the incoming stimulus, may cause fMRI activity patterns tuned to probability. In this view, had the probabilities not been frequently updated, we may not have detected them in activity patterns. This possibility remains to be tested.

We now turn to discuss some limitations of our approach. First, we have used binary sequences with stimulus A or B, so that the generative process can be described equivalently in terms of p(A) or p(B), since p(A)=1-p(B). We found evidence that some pieces of the cortex exhibit similar activity for values of p(A) that are symmetric with respect to 0.5, e.g. when p(A)=0.2 and p(A)=0.8 (see Supplementary Fig. 4). This may be due to a representation of the event probability switching between p(A) and p(B), or due to a similar proportion within a voxel of neurons each coding for either p(A) or p(B). Another possibility is that some representations of probability may focus on the probability of the most likely stimulus (the one with p>0.5) and the identity of this stimulus (these two pieces of information are sufficient to reconstruct p(A) and p(B) in the binary case). The use of more than 2 items in a sequence would be useful to adjudicate between these different possibilities.

Invariance is a useful criterion for testing for neural representations, especially in the context of uncertainty (Walker et al., 2023). The invariance of probability representation could be tested with respect to the features of the stimulus (e.g. visual stimuli that differ in shape, or color, rather than orientation) or the sensory modality used (e.g. auditory stimulus (Meyniel & Dehaene, 2017)). The origin of the probability could also be changed. Here, the probability originates from a statistical learning process operating on a sequence, but not all probabilities do. Reasoning and memory can also be used, for example when estimating the probability that a statement is true, such as “Is Paris bigger than Berlin?” (Lebreton et al., 2015).

In addition, the timing of the task and the poor temporal resolution of fMRI precluded the use of trial-level decoding, which was done at the session level here. Future studies could explore a decoding of probability across trials, perhaps benefiting from the use of better time-resolved recordings such as electrophysiology.

In summary, we have unraveled a neural representation of event probability in the human cortex that is based on a highly nonlinear code, and that cannot be detected by simpler methods that assume a linear code. The methods we have developed here can be used to search for neural representations of probability in many different types of tasks.

## Methods

### Participants and task

The behavioral and fMRI data used in this paper have already been analyzed in (Bounmy et al., 2023). Experimental protocols were approved by the local ethics committee (CPP-100032 and CPP-100055 Ile-de-France) and the informed consent of all 29 participants (15 female, mean age 25.4 ±1.0 s.e.m.) was obtained before they began the experiment. Three participants were excluded from analysis due to acquisition problems, resulting in an effective total of N=26 subjects for analysis.

After receiving task instructions and completing a training session, participants entered the MRI scanner and performed 4 task sessions, during which the scanner recorded functional MRI data. Each session lasted approximately 11 minutes and consisted of a sequence of 420 stimuli presented one after the other, for 1 s each, with an inter-stimulus interval of 0.3 s (see example in Fig. 1A).

Each stimulus had a binary value, A or B, represented in the task by one of two distinct orientations of a high-contrast grating, easily distinguishable from each other. The values were drawn from a Bernoulli distribution with a hidden generative probability p(A) to be learned, whose value will be denoted *h_t_*. The hidden probability *h_t_*followed a stochastic change-point process: at each time step, it could either remain the same or undergo a change point, with a change-point probability of 1/75 under the constraint of a maximum period of 300 stimuli without change points. The value of *h_t_* was uniformly sampled between 0.1 and 0.9 initially and at each change point, under the constraint that the odds of A changed by a factor of at least four.

Throughout the task, the subjects’ goal was to estimate the hidden probability *h_t_*. The occurrence of unsignaled and unpredictable change points made the estimation all the more challenging. Accurate estimation requires making probabilistic inferences about the value of *h_t_* given the observed stimuli, with knowledge of the generative process of the sequences, as described in the task model below. Subjects could make such inferences as they had been briefed about the generative process in an informal way during the instructions phase.

Subjects were instructed to continuously estimate the probability during the sequence, and were occasionally asked to report their estimate during dedicated time periods. Isolating the report periods from the periods of stimuli and the estimate updates they elicited allowed us to ensure that neural representation of the factors of interest we found in the fMRI recordings, related to the estimation process, were not confounded by the subject’s reporting. Report periods occurred on average every 22 stimuli, with a random uniform jitter of ±3 stimuli maximum. Each report period consisted of a response screen where the subject made two choices: a choice of estimate range for *h_t_* among three or five possible ranges (the scale was randomly selected between the three-choice scale and the five-choice scale for each period, except for the first half of participants where the scale was always the five-choice scale), and a choice of confidence level associated with this estimate among five possible levels. These choices were made using a right-handed five-finger button pad. Although the choices were discrete, subjects were instructed to internally generate continuous-valued estimates. They were further incited to do so in order to produce correct reports as they did not know in advance which scale, and therefore which choice ranges would be available (there being no trivial mapping between the two scales).

### Task model

The model we used for the task is the normative model, in the sense that it produces, for each trial, the optimal estimate of the hidden probability that the subject could have produced, given the stimuli they have observed so far and their knowledge of the generative process of the task. This estimation problem requires inferring the value of *h_t_*given the stimulus values observed in the past *s_1:t_*. Since the generative process is probabilistic, the inference problem is also probabilistic (the value of *h_t_* cannot be determined with certainty). Using the rules of probability calculus and in particular Bayes’ rule, a posterior distribution on *h_t_* can be calculated, denoted as *p(h_t_ | s_1:t_)*.

In the present case, one solution to calculate algorithmically the posterior distribution is to proceed iteratively on the observations, initializing *p(h_0_)* to a uniform distribution and computing *p(h_t+1_ | s_1:t+1_)* based on the value of *s_t+1_* and the previously computed *p(h_t_ | s_1:t_)*. This calculation is done using the following formula.

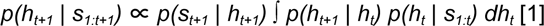

This formula is derived from the rules of probability calculus by leveraging two conditional independence properties of the present generative process: (1) *s_t+1_*is conditionally independent of *s_1:t_* given *h_t+1_*; (2) *h_t+1_* is conditionally independent of *s_1:t_* given *h_t_*. See Supplementary Note 1 for the derivation.

The ∝ operator denotes the equality up to a constant factor. This constant is implicitly factored in at computation time by normalizing the right-hand side of equation [1] so that it sums up to 1 over the possible values of *h_t+1_*, to obtain the left-hand side. The term *p(s_t+1_ | h_t+1_)* is given by the Bernoulli distribution, equal to *h_t+1_* or *(1-h_t+1_)* depending on whether s_t+1_ is A or B. The term *p(h_t+1_ | h_t_)* reflects the generative probability that a change point occurs (1/75) or does not occur (74/75), depending on whether *h_t+1_*is different from or equal to *h_t_*, respectively.

The normative estimate of hidden probability and the associated confidence are both calculated from the posterior distribution. The probability estimate is equal to the mean of the posterior, *E[h_t+1_ | s_1:t_]*. It is optimal in the sense that it minimizes the mean squared error with the true value, and is equal to the posterior probability that the next stimulus value is A (this is formally what subjects were asked to report). Confidence was defined as *-log SD[h_t+1_ | s_1:t_]*, the log-precision of the posterior (up to a factor of two). Hereafter, we will use the symbol *x* to refer to these two types of estimates indiscriminately (*x_t_*representing the estimate for stimulus *s_t_*) as the presented encoding models are mostly independent of the specific type of estimate being encoded.

The task model was implemented using Python and the NumPy package (https://numpy.org).

### Encoding models

Encoding models predict the fMRI activity of a voxel as a function of the estimates obtained from the stimulus sequence seen by the subject. We defined four main encoding models, 2×2, depending on whether the encoded estimates are probability or confidence estimates, and whether the model tuning curve function belongs to the linear or the versatile class (Fig. 2).

We also considered another model, following a hypothesis proposed in the literature, in which the activity was a function of the entire posterior distribution (rather than a moment of the distribution, like the probability and confidence estimates are) (Sahani & Dayan, 2003; Zemel et al., 1998). That is, in that model, *f_i_(x)* in equation [3] was replaced by its posterior mean ∫*f_i_(x)p(x)dx*, where *p* is the posterior distribution. However, when we tested that model on our fMRI data, it explained the data less well than the simpler models encoding the probability or confidence estimates (see Supplementary Fig. 3). Therefore, we focused on the simpler models.

The probability and confidence estimates are calculated from the sequence as explained in the “Task Model” section above. The model tuning curve function maps the encoded estimate, *x,* to a prediction of neural activity, *ŷ*, and is parameterized with weights *w* that are to be fitted to the data. In the linear class, this function is of the form:

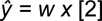

In the versatile class, this function is of the form:

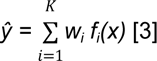

where the *f_i_* are (radial) basis functions that have approximation properties (Franke, 1982). Here, we used Gaussian basis functions, but we also performed simulations using sigmoid basis functions, and the results were similar whether we used Gaussian or sigmoid functions (Supplementary Fig. 2). The Gaussian basis functions are expressed *f_i_(x) = c exp[-(x-*𝜇*_i_)^2^ / (2*𝜎*^2^)]*. The centers of the basis functions *µ_i_* were distributed to have equal spacing between two consecutive centers, between the lower bound of the interval and the first center, and between the upper bound and the last center (the interval being [0, 1] for probability, and [1.1, 2.6] for confidence). The number of basis functions was *K*=10, and their dispersion was 𝜎=0.04 for probability and 𝜎=0.06 for confidence. These values were determined through simulations to optimize the *R^2^* scores averaged over a wide range of simulated activity with different *K* and 𝜎 values.

To transform the predictions of the theoretical models described above into predictions at the level of fMRI activity in a voxel, we convolved the encoding model regressors (that is, the scalar quantities that are multiplied with the weights *w: x* in the linear case and the *f_i_(x)* in the versatile case), with the canonical hemodynamic response function at the onsets of the corresponding stimuli (these convolved regressors are often referred to as “parametric modulations” in the fMRI community).

Additionally, we included in the encoding model other regressors corresponding to factors of no-interest in this study, which we removed during testing to evaluate the specific encoding of probability or confidence. The regressors of no-interest were the following.

● Parametric modulations of stimulus onsets associated with factors other than probability and confidence:

○ a constant
○ the Shannon surprise induced by the stimulus, *-log p(s)*, where *s* is the stimulus value and *p(s)* is the normative probability estimate for that stimulus value given the previously observed stimuli
○ the Shannon entropy of the outcome implied the normative probability estimate, *-p log(p) - (1-p) log(1-p)* (see Supplementary Fig. 4).
● Parametric modulations of response screen onsets modeling reporting periods (including, for each period, a response screen for probability and another for confidence):

○ a constant
○ the normative estimate
○ the estimate reported by the subject
● Six motion regressors

Finally, we applied the same temporal preprocessing to all regressors as we applied to the fMRI signal (detrending, filtering, z-scoring across sessions, and session-wise demeaning, see MRI data preprocessing section).

At the voxel level, the fMRI activity predicted by the model after preprocessing, *ŷ_fMRI_*, is written as *ŷ_fMRI_ = w ẋ + **w_n_ n*** for the linear model, and *ŷ_fMRI_ = **w ḟ**(x) + **w_n_ n*** for the versatile model, where · represents the convolution and temporal preprocessing operations, ***w*** and ***f*** are the vectors *[w_1_, …, w_k_]* and *[f_1_, …, f_k_], **n*** is the vector comprising all preprocessed regressors of no-interest, and **w_n_**is their associated weights vector (bold symbols denote vectors as opposed to scalar quantities).

The encoding models were implemented using Python, NumPy (https://numpy.org), and nilearn (https://nilearn.github.io).

### Evaluation of the encoding models

We evaluated the ability of the encoding models to predict fMRI data for a given subject using leave-one-session-out cross-validation: three sessions were used for training the model, and the fourth, left-out session, was used to test the trained model. This procedure was repeated for each of the four possible choices of the left-out session, and the test scores obtained were averaged across the four left-out sessions.

#### Training

During training, the weights of the encoding model were fitted using Ridge regression. The Ridge penalty was calculated using an analytical formula that adjusted the amount of penalty (𝜆) based on the number of model parameters (*m*): 𝜆*=199m*, which was validated through simulations.

#### Testing

During testing, we calculated the fMRI activity predicted by the model using only the part of the model associated with the factors of interest (probability or confidence). (This is equivalent to replacing the regressors of no-interest with their mean value for the session, which is equal to 0 after session-wise demeaning.) As a score, we first calculated the *R^2^* obtained by comparing the model predictions with the actual fMRI data, using the sums-of-squares formulation (Poldrack et al., 2020). We then calculated a null distribution of *R^2^* scores by injecting null predictions into the *R^2^* calculation. These null predictions were obtained by replacing the true stimulus sequence seen by the subject with another sequence randomly generated according to the task generative process, for 100 generated sequences. Finally, we calculated the score we call *z-R^2^* by standardizing the *R^2^* score obtained for the true sequence by the mean, *µ_0_(R^2^)*, and standard deviation, 𝜎*_0_(R^2^)*, of the null distribution: *z-R^2^ = (R^2^ - µ_0_(R^2^)) /* 𝜎*_0_(R^2^)*.

Models were fitted and tested using Scikit-Learn (https://scikit-learn.org).

### Simulation of encoding models

We conducted simulations to verify that our procedure, applied in our experimental protocol, was able to detect and differentiate a neural encoding of probabilities or confidence, linear or nonlinear. For this purpose, experimental data was generated assuming a certain encoding model, and the generated experimental data was analyzed with other encoding models. As for subjects, each generated experiment consisted of four sessions, with one sequence of stimuli per session, generated according to the task process. Noisy fMRI activities were then generated for each session and a certain number of simulated voxels assuming a certain generative model of neural activity, which corresponded to one of the four encoding models presented above (encoding of probability or confidence estimates, according to a linear or a versatile encoding model).

The procedure for generating fMRI activity was as follows. For each simulated voxel, we randomly generated weights for the generative model by drawing each weight uniformly in [-0.5, 0.5]. We generated the “signal” component of the fMRI activity in accordance with the generative model, by calculating the weighted sum of the corresponding fMRI regressors as described in the above section on encoding models. We then injected Gaussian white noise with power equal to nine times that of the signal, in order to obtain a signal-to-noise ratio of 10% signal to 90% noise. This produced a set of fMRI activities for each generated experiment and each possible generative model.

For each generated experimental data, the procedure presented in the above section was used to fit the encoding models and evaluate their ability to predict the simulated fMRI data (as subsequently done for the subjects’ fMRI data). This produced one *z-R^2^* score per voxel, fitted model, generative model, and generated experiment. We averaged the scores obtained across one hundred experiments and one hundred simulated voxels to obtain an average *z-R^2^* score for each possible generative model × fitted model pair, resulting in a 4×4 matrix, shown in Fig. 2B.

This analysis was also performed by splitting the versatile model into one with Gaussian basis functions and one with sigmoid basis functions to produce the matrix shown in Supplementary Fig. 2.

The simulations were implemented using Python and Numpy.

### MRI data acquisition

#### Equipment

The MRI scanner was a Siemens MAGNETOM 7 Tesla at the NeuroSpin center (CEA Saclay, France), with whole-body gradient and 32-channel head coil by Nova Medical.

#### Functional MRI acquisition

Whole-brain functional volumes with 1.5mm isotropic voxels and T2*-weighted fat-saturation images were acquired using a multi-band accelerated echo-planar imaging sequence (Moeller et al., 2010; https://www.cmrr.umn.edu/multiband/). The sequence parameters were: multi-band factor = 2 (MB); GRAPPA acceleration factor = 2 (IPAT); partial Fourier = 7/8 (PF); matrix = 130 × 130, number of slices = 68, slice thickness = 1.5 mm, repetition time = 2 s (TR); echo time = 22 ms (TE); echo spacing = 0.71 ms (ES); flip angle = 68° (FA); bandwidth = 1832 Hz/px (BW); phase-encoding direction: anterior to posterior.

Before each session, two single-band functional volumes were acquired with the above parameters except that they had opposite phase-encoding directions. This was later used for distortion correction (see below in *Preprocessing*).

A Gradient Recalled Echo (GRE) sequence was acquired for calibration before starting the sessions. For shimming, a B_0_ map was acquired and loaded in the console, first of the whole brain, then in an interactive fashion on the occipito-parietal cortex. This aimed to reduce the FWHM and increase the T2*. A B_1_ map was then acquired and the intraparietal sulcus values were used to compute a reference voltage from which the system voltage was chosen.

#### Anatomical MRI acquisition

Whole-brain T1-weighted anatomical images with 0.75 mm isotropic resolution were acquired using an MP2RAGE sequence. The sequence parameters were: GRAPPA acceleration factor = 3 (IPAT), partial Fourier = 6/8 (PF), matrix = 281 × 300, repetition time = 6 s (TR), echo time = 2.96 ms (TE), first inversion time = 800 ms (TI_1_), second inversion time = 2700 ms (TI_2_), flip angle 1 = 4° (FA_1_), flip angle 2 = 5° (FA_2_), bandwidth = 240 Hz/px (BW).

### MRI data preprocessing

The functional volume slices were corrected for slice-timing with respect to the slice acquired in the temporal middle of a volume acquisition. The functional volumes were corrected for motion using rigid transformations, and co-registered to the anatomical image. Session-wise distortion correction was applied to the functional volumes using FSL apply_topup, after having estimated a set of field coefficients for the session using FSL TOPUP with the two-single band volumes with opposite phase-encoding directions acquired before that session.

Prior to encoding and decoding analyses, the fMRI time series of each voxel and session were temporally detrended and high-pass filtered at a cutoff frequency of 1/128 Hz. The series from the four sessions were then concatenated in order to z-score the fMRI data across the four sessions for each voxel for numerical convenience. Note that we did not z-score the fMRI data per session because under the versatile encoding model, it is expected that the variance of the fMRI signal should change from session to session depending on the probabilities and confidence levels estimated during each session. Finally, the mean of each session was subtracted from the data in order to ignore any changes in the signal baseline between sessions that might be caused by nuisance factors.

Slice-timing correction, motion correction, and co-registration were done using tools from SPM12 (https://www.fil.ion.ucl.ac.uk/spm/software/spm12). Distortion correction was done using tools from FSL (https://fsl.fmrib.ox.ac.uk/fsl/fslwiki/FSL). SPM12 and FSL were called from Python using the NiPype module (https://nipype.readthedocs.io). Further preprocessing was done using Python and the NumPy, SciPy (https://scipy.org) and nilearn (https://nilearn.github.io) packages.

### Conversion of volumetric data into cortical surface data

Here we detail the processing steps we used throughout this study to project subject-level volumetric data, such as the *R^2^* and *z-R^2^* scores computed from the fMRI data (see section on encoding model evaluation), onto the cortical surface, and to bring them into a common space across subjects. These processing steps were performed using FreeSurfer (https://surfer.nmr.mgh.harvard.edu) and the Python interface to FreeSurfer commands provided by the NiPype module. The resulting surface data were then analyzed by working directly with the numerical arrays in Python, unless otherwise stated.

For each subject, the cortical surface was reconstructed from the acquired high-resolution anatomical MRI image by running the FreeSurfer command ‘recon-all’. These reconstruction data were then used to project other volumetric data calculated from the functional data onto the cortical surface, working in the subject’s native space. This projection was performed using the ‘mri_vol2surf’ command. The surface data were then normalized, i.e., resampled to be brought into a common space: the ‘fsaverage’ template of FreeSurfer, version 7 high-resolution (163,842 vertices per hemisphere), and were spatially smoothed with a Gaussian kernel of 3mm full width at half maximum. The resulting data are surface maps containing one data point per vertex (a vertex is the surface equivalent of a voxel in the volume, called vertex because vertices are assembled to define polygons that together form a three-dimensional mesh of the cortical surface).

### Group-level statistical maps

The subjects’ *z-R^2^* surface maps were grouped together and a one-sample t-test against zero was performed at the group level. The resulting p-value maps were thresholded at the vertex level with a threshold of p<0.001, and corrected for multiple comparisons with a family-wise error rate of p_FWE_<0.05 using FreeSurfer’s Monte Carlo simulation-based cluster correction with a vertex-wise cluster-forming threshold of p<0.001.

### Definition of vertices of interest for characterization

To ensure the reliability of the tuning curves used for characterization, we defined a set of vertices of interest for which the encoding signal was sufficiently strong. One set was defined per type of estimate and per subject, as the vertices with the strongest encoding signal are not the same for probability and confidence, and vary across subjects. Additionally, the number of vertices was adjusted depending on the availability of vertices with a strong enough signal.

The definition was done in two steps: first at the group level, and then at the subject level. At the group level, a region of interest was defined as the union of parcels from the HCP-MMP1.0 atlas (Glasser et al., 2016) containing the significant clusters found in the group-level statistical maps obtained for the corresponding estimate type (clusters shown on Fig. 3 and 4 and listed in Table 1 and Supplementary Table 1). The vertices of interest for each subject were then defined within the group-level region, using the following steps. The *R^2^* scores obtained with the encoding model were converted into p-values in each subject and vertex. For each vertex and subject, a p-value was calculated from the *R^2^* score as the probability of obtaining a value at least as large in a distribution of *R^2^* values obtained when the fMRI data is replaced with white noise (empirically calculated for each subject with 1,000,000 noise samples). Finally, the vertices with FDR-corrected p<0.05 were selected as vertices of interest (controlling the False Discovery Rate using the Benjamini-Hochberg procedure)).

As mentioned in the Results section, we verified the reliability of our estimations within the defined vertices of interest using a test-retest procedure, by fitting the weights of the versatile encoding model on two independent halves of the data and comparing the estimated weights between the two halves.

### Characterization of tuning curves

Three characteristic measures were defined and applied to the tuning curves estimated for each vertex of interest.

*1) Proportion of tuning curves maximized at non-extreme values.* For each tuning curve, we took the input value at which the curve was maximized (i.e. the argmax of the tuning curve function) and treated it as non-extreme if it fell between the lower and upper bounds of the domain with a margin of at least twenty percent relative to the span of the domain. The proportion refers to the proportion of vertices out of all vertices of interest within the subject.
*2) Non-monotonicity index.* Mathematically speaking, a differentiable function is said to be monotonic if its derivative remains of the same sign over its domain. By extension, we defined the monotonicity index *m(f)* of a function as the absolute value of its average derivative, normalized such that the index of any purely monotonic function is equal to 1. The non-monotonicity index *n(f)* was *1 - m(f)*. It is calculated by the formula *n(f) = 1 - |f(x_max_) - f(x_min_)| / (f_max_ - f_min_)*, where *x_min_* and *x_max_* are the lower and upper bounds of the domain, and *f_min_* and *f_max_* are the minimum and maximum of the function over the domain, respectively. The indices calculated for each tuning curve function were averaged across the vertices of interest to obtain a single value per subject. Note that *m(f)*=1 is a necessary condition for a monotonic function, but not a sufficient one (non-monotonic functions can yield *m(f*)=1).
*3) Nonlinearity index.* We defined the nonlinearity index at the vertex level as the degree of the polynomial that best fits the estimated tuning curve for that vertex. To avoid favoring larger degrees due to overfitting, we used the tuning curves estimated on two independent halves of the data, one to fit the polynomial model for each degree, the other one to measure the variance explained (*R^2^*) by the fitted polynomials. The nonlinearity index was equal to the degree of the polynomial that led to the best *R^2^* score. To summarize the computed index values at the subject level, we took the median across vertices. At the group level, we tested statistical significance using a Kruskal-Wallis test by ranks (a nonparametric method for testing whether samples originate from the same distribution).

### Decoding of probability and confidence

The same method was used for decoding probability and confidence; here we explain the method in the case of probability. Decoding was performed at the level of each session and subject, and it leveraged the methods used for the encoding analysis. The encoding model characterized a pattern of voxel responses to (overlapping) bins of probability as a set of regression weights assigned to Gaussian basis functions. The decoder aimed to identify the probability bin corresponding to the response pattern measured across several voxels in a test session. Decoding accuracy was assessed with leave-one-session-out cross-validation. The encoding model was fitted (with 5 basis functions instead of 10) separately in the training set (three sessions) and the left-out session. The decoder assigned a probability bin to a given response pattern on the test set by identifying the probability bin eliciting the most similar response pattern in the training set. More precisely, the decoder used the correlation distance *D*(*j*, *k*) between the response patterns corresponding to the *j*-th probability bin in the test session and the *k*-th bin in the training set, and looked for the *k* that minimized *D* for a given *j*. The correlation distance is one minus the Pearson correlation (Walther et al., 2016) qualitatively similar results, although inferior, were obtained with the Euclidean distance.

The decoder was applied to selected voxels in each parcel of the Glasser et al atlas (Glasser et al., 2016). This atlas is available in the fsaverage template used for anatomical normalization (https://figshare.com/articles/HCP-MMP1_0_projected_on_fsaverage/3498446). The atlas was projected onto the native anatomical surface of each subject using the inverse normalization transform, and projected back into their native volume, both with FreeSurfer. The voxels corresponding to each parcel in the functional images were identified based on the coregistration of functional and anatomical images. Decoding was applied to the 100 voxels with the largest *z-R²* value, estimated within the training sessions only.

For each subject, decoding accuracy was assessed for each probability bin as the fraction of correctly assigned probability bins (chance level is 1 out of 5 possibilities) on each left-out session, and averaged across the left-out sessions. Some probability bins concerned fewer than 5% of stimuli in some test sessions, which we deemed unreliable; the corresponding session-level accuracy was omitted from the average across sessions. More precisely, we considered that a stimulus fell in a given bin if its probability elicited more than 10% of the maximum value of the corresponding basis function. The significance of decoding accuracy was assessed, for each parcel, with a two-sided t-test against chance-level accuracy at the group level, and corrected for multiple comparisons across parcels with false discovery rate (0.05) correction (Benjamini-Hochberg procedure).

### Analysis of Representational Dissimilarity Matrices (RDMs)

The same method was used for probability and confidence; here we explain the method in the case of probability. We analyzed the distance matrices *D*(*j*, *k*) used for the decoding process, which are also known as representational dissimilarity matrices (RDM). For each subject the RDM of each parcel was the average RDM obtained across cross-validation folds (ignoring the rows *j* of each RDM corresponding to probability bins concerning fewer than 5% of stimuli in a session). Note that since RDMs were estimated with leave-one-session-out cross-validation, they are not symmetric and their diagonal is not 0. The empirical RDMs were compared to the RDMs of different neural codes, namely, the identity RDM that should arise from a highly nonlinear code, and the graded RDM that should arise from a linear (and more generally monotonic) code. We estimated the regression weights corresponding to each theoretical RDM in a multiple regression analysis; theoretical RDMs were z-scored to make the regression weights commensurable.

## Data availability

Behavioral data, raw and preprocessed MRI data will be made available on a public repository upon acceptance.

## Code availability

All analysis code to reproduce the reported results and figures from the shared data will be made available on GitHub upon acceptance.

## Acknowledgements

This work was supported by funding from the French National Research Agency (ANR grand #18-CE37-0010-01) and from the European Research Council (ERC grant #947105) to F. M. C.F. was supported by a PhD fellowship from ENS Paris-Saclay (France).

## Author contributions

CF: Conceptualization, Methodology, Software, Validation, Formal analysis, Writing—original draft, Writing—review & editing, Visualization; TB: Conceptualization, Methodology, Software, Investigation; SD: Methodology, Software; BT: Conceptualization, Methodology, Writing—review & editing; EE: Conceptualization, Methodology, Writing—review & editing; FM: Conceptualization, Methodology, Software, Formal analysis, Resources, Writing—original draft, Writing—review and editing, Visualization, Supervision, Project administration, Funding acquisition.

## Supplementary information

### Supplementary Notes

#### Supplementary Note 1

Equation to be demonstrated:

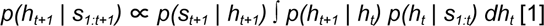

Below is the mathematical proof of the formula given by equation [1].

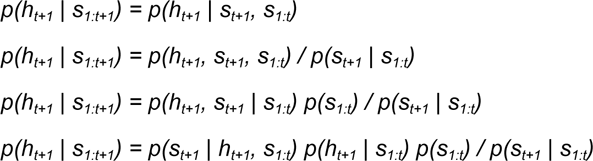

Conditional independence property (1): *s_t+1_* is conditionally independent of *s_1:t_* given *h_t+1_*, therefore:

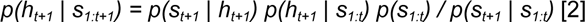

The first term of equation [2] is the same as that equation [1]. Let’s expand the second term of equation [2] using the sum rule.

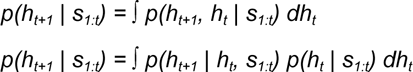

Conditional independence property (2) : *h_t+1_* is conditionally independent of *s_1:t_* given h_t_, therefore

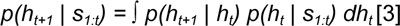

By combining equations [2] and [3] and omitting the normalization factor *p(s_1:t_) / p(s_t+1_ | s_1:t_)*, we obtain equation [1] which was to be demonstrated.

### Supplementary Figures

**Supplementary Figure 1.**
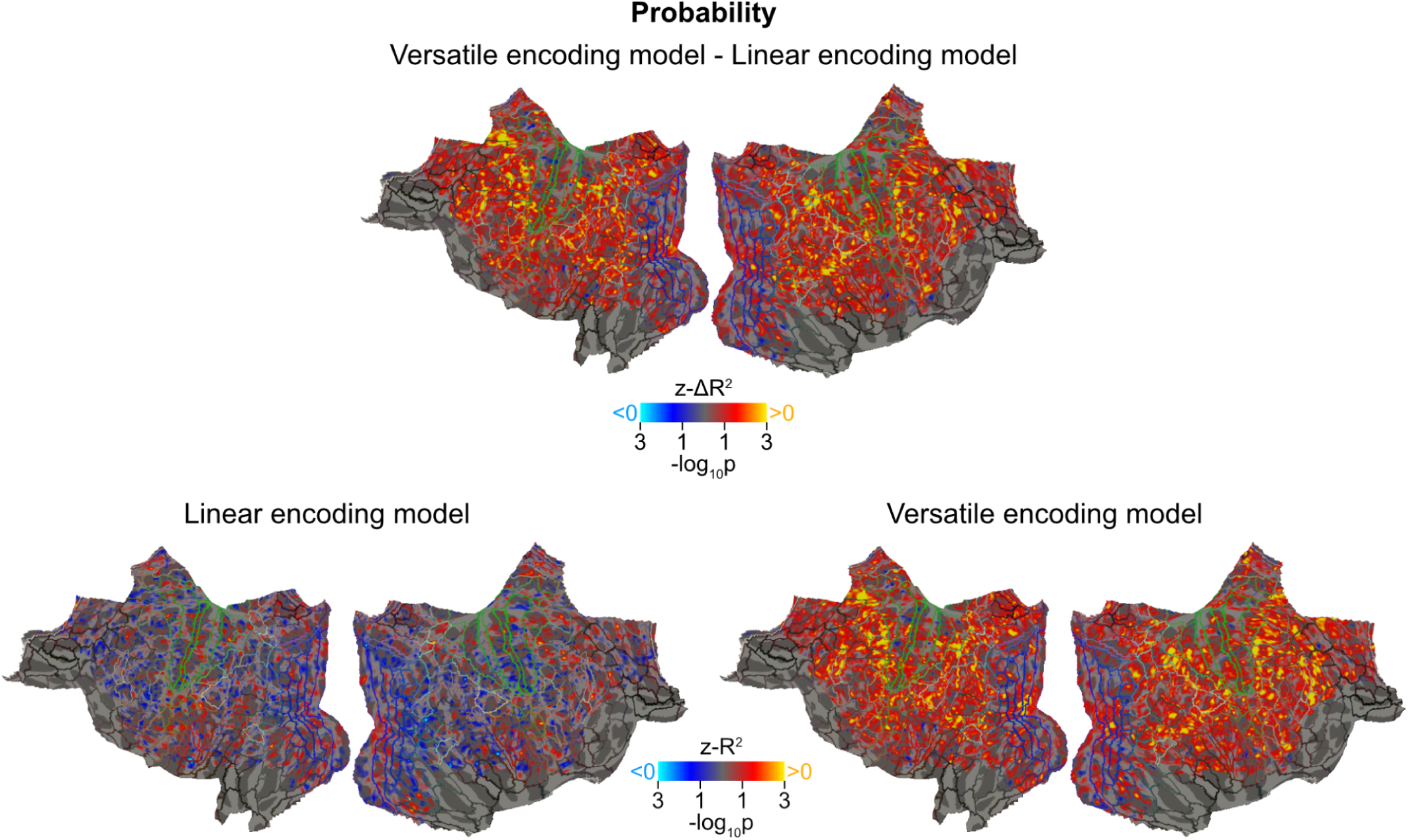
Difference in cross-validated variance explained by the versatile and the linear encoding models for probability. Vertex-wise p-values are shown on flattened maps of the cortex. Top: P-values correspond to the group-level significance of z-ΔR^2^ scores obtained across subjects (ΔR^2^ = R^2^[versatile] - R^2^[linear], cold/hot colors favor the linear/versatile encoding model respectively, p values are unthresholded and uncorrected). Bottom: Maps of the individual models, repeated from Fig. 3 for illustration purposes. Delineated by colored lines is the HCP-MMP1.0 parcellation (Glasser et al., 2016).

**Supplementary Figure 2.**
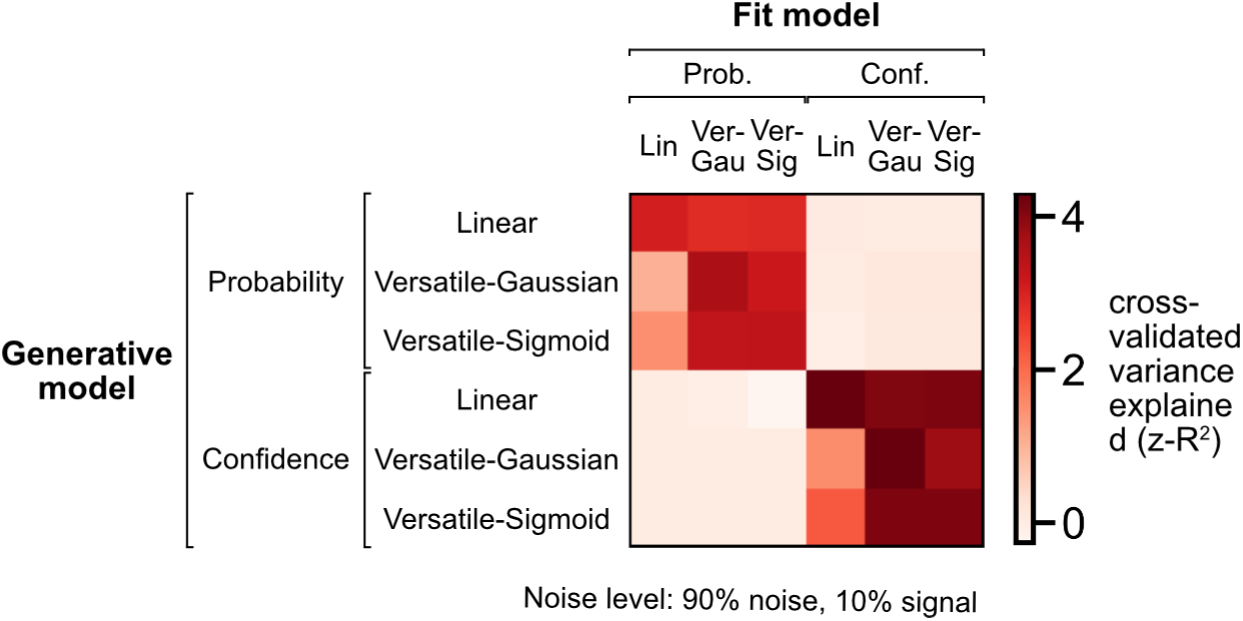
Simulation results with Gaussian and sigmoid basis functions. Simulation results obtained as in Fig. 2B after splitting the versatile encoding model into two: one with Gaussian basis functions (the one used in the main text, referred to as Versatile-Gaussian above) and one with sigmoid basis functions (referred to as Versatile-Sigmoid above). The sigmoid basis functions of the Versatile-Sigmoid model are expressed *f_i_(x) = 1 / [1 + exp[-k(x-*𝜇*_i_)]].* For comparison, we took the same number of basis functions (10) and the same centers 𝜇*_i_*as for Versatile-Gaussian, and an equivalent slope *k* was computed from the Versatile-Gaussian’s 𝜎 using the formula *k = 4 / [(2π)^1/2^*𝜎].

**Supplementary Figure 3.**
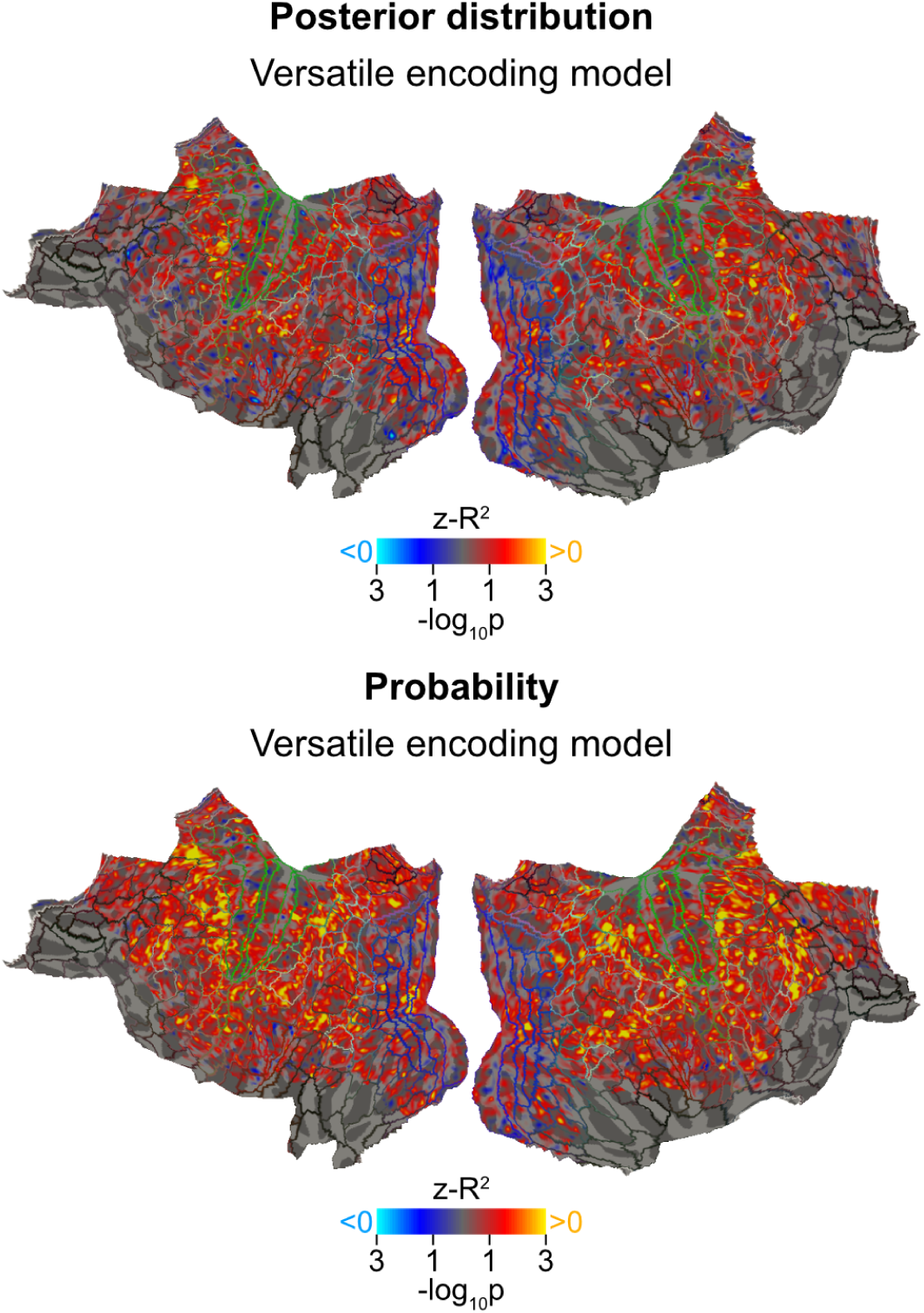
Variance explained by the model encoding the posterior distribution (top) vs. the probability estimate (bottom). Cortical maps as in Fig. 3. See Methods for the encoding model of the posterior distribution. Bottom map is repeated from Fig. 3 for illustration purposes.

**Supplementary Figure 4.**
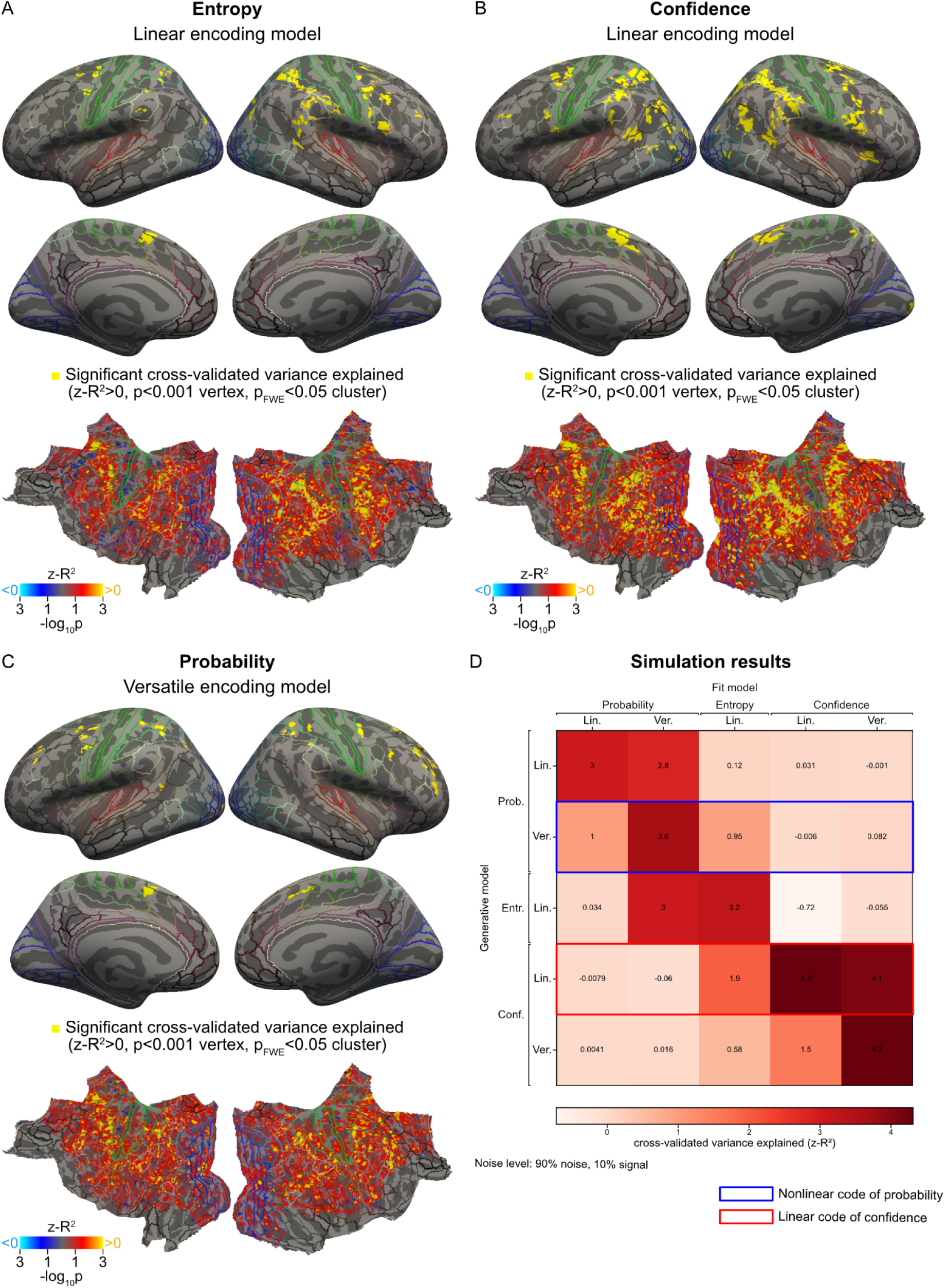
The (linear) effect of confidence and the (nonlinear) effect of probability are not confounded by a (linear) effect of entropy. (A) Cortical maps for the linear model of entropy, which quantifies the extent to which the next stimulus is unpredictable (entropy is maximum for p(A)=0.5 and gradually decreases as p(A) deviates from 0.5). Plotting conventions are as in Fig. 3. Panels B and C for confidence and probability are reproduced from Fig. 4 and Fig. 3, respectively, to facilitate comparison with A. Note that the regression models in B and C include entropy as an additional regressor, but that confidence and probability are not included as additional regressors in A. The effect of entropy seen in A is less widespread than the effect of confidence, and largely overlaps with the latter; the effect seen in A is thus likely to arise from the small correlation that exists between entropy and confidence. The effect of confidence we observed in B cannot be reduced to an effect of entropy because the latter effect is included in the regression model, and the effect of confidence is more widespread than the effect of entropy. The (nonlinear) effect of probability in C is also not confounded by a (linear) effect of entropy since entropy is included as an additional regressor, and since the effects of probably and entropy are anatomically distinct (at least in the anterior part of the dorsolateral prefrontal cortex). (D) Simulation results. Plotting conventions are as in Fig. 1B. Note that a versatile model of probability explains the data generated by a linear model of entropy very well, but the converse is not true. Also note that a linear model of entropy can only partially explain the data generated by a linear model of confidence. The pattern of results observed in the data (panels A-C) are consistent with the results obtained by simulations.

**Supplementary Table 1.**
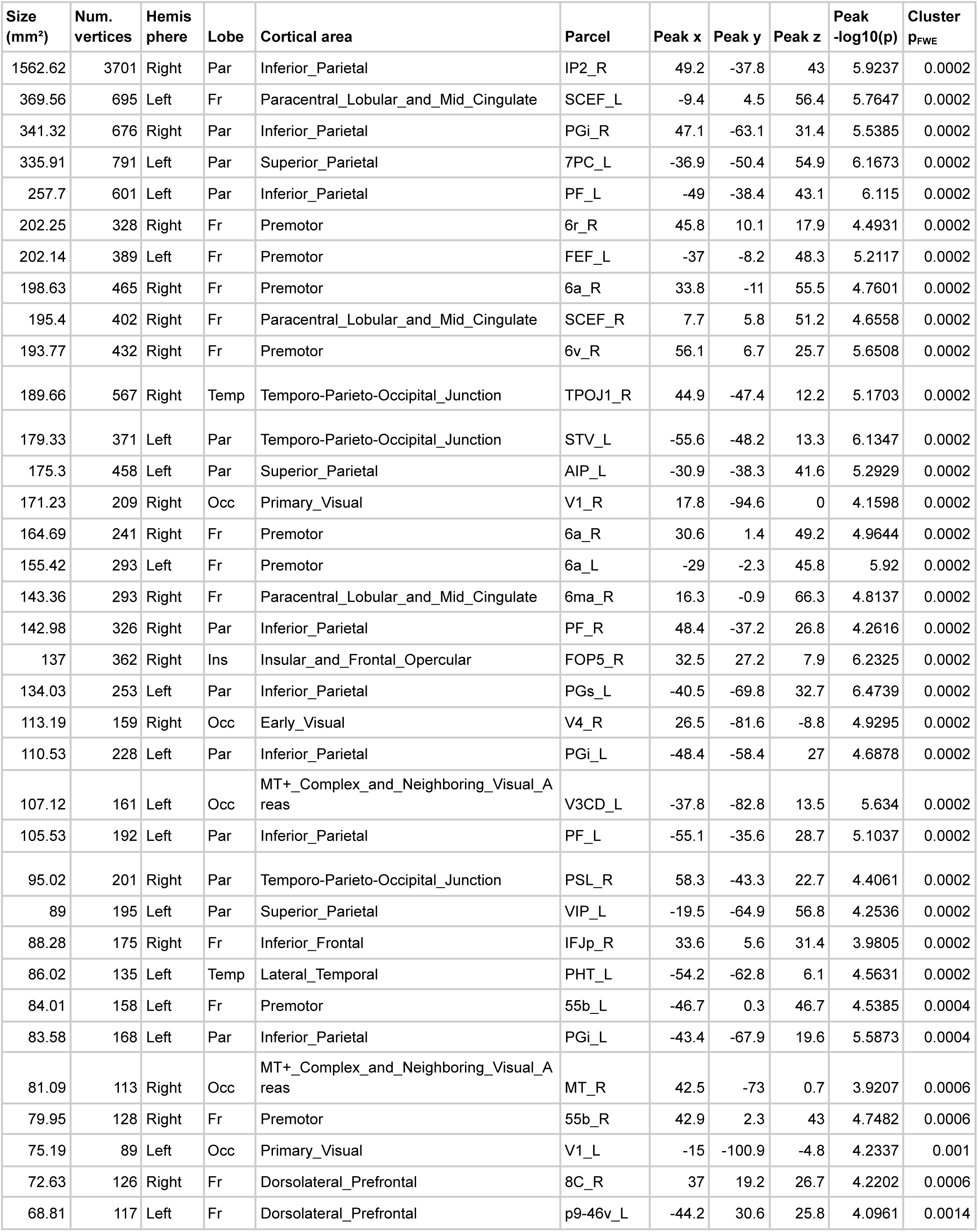
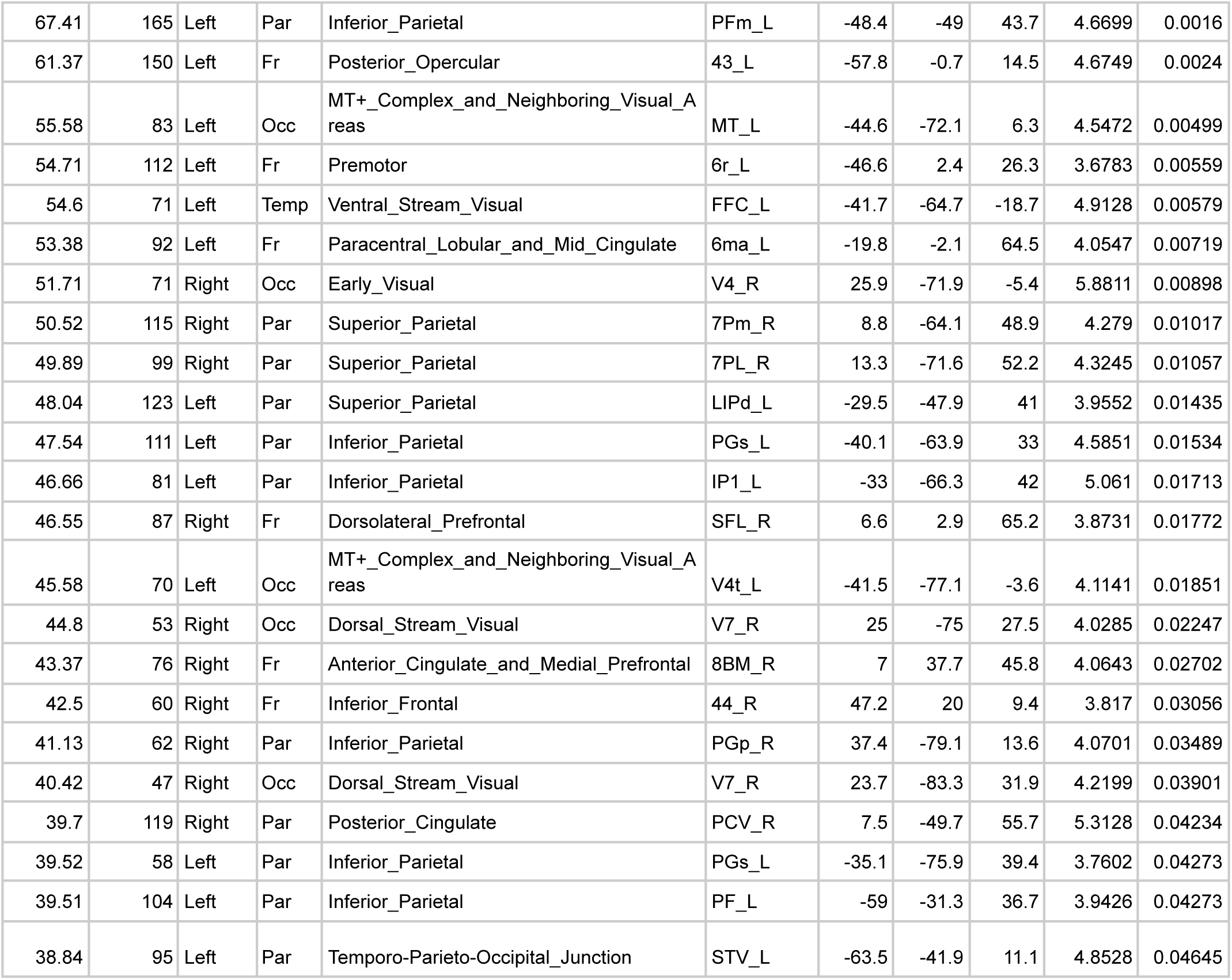
Significant clusters explained by the linear encoding model for confidence. The names of the cortical areas and parcels refer to the HCP-MMP1.0 atlas (Glasser et al., 2016). Peak x, y, z are MNI coordinates.

